# Suicidal chemotaxis in bacteria

**DOI:** 10.1101/2021.12.21.473623

**Authors:** Nuno M. Oliveira, James H. R. Wheeler, Cyril Deroy, Sean C. Booth, Edmond J. Walsh, William M. Durham, Kevin R. Foster

**Affiliations:** Department of Zoology, University of Oxford, Oxford, UK; Department of Applied Mathematics and Theoretical Physics, University of Cambridge, Cambridge, UK; Department of Veterinary Medicine, University of Cambridge, Cambridge, UK; Department of Biochemistry, University of Oxford, Oxford, UK; Department of Physics and Astronomy, University of Sheffield, Sheffield, UK; Department of Engineering Science, University of Oxford, Oxford, UK

**Author notes:** These authors contributed equally (Nuno M. Oliveira, James H. R. Wheeler). Email: Correspondence and requests for materials should be addressed to William M. Durham or to Kevin R. Foster. **Author Contributions:** N.M.O., J.H.R.W., C. D., E.J.W., W.M.D. and K.R.F. designed research; N.M.O., J.H.R.W., C.D. and S.C.B. performed research; N.M.O. and J.H.R.W. and W.M.D. analyzed data; N.M.O., J.H.R.W., W.M.D. and K.R.F. wrote the paper.

**Keywords:** bacterial biofilms, antibiotic gradients, chemotaxis, bacterial suicide, competition sensing

## Abstract

Bacteria commonly live in communities on surfaces where steep gradients of antibiotics and other chemical compounds routinely occur. While many species of bacteria can move on surfaces, we know surprisingly little about how such antibiotic gradients affect cell motility. Here we study the behaviour of the opportunistic pathogen *Pseudomonas aeruginosa* in stable spatial gradients of a range of antibiotics by tracking thousands of cells in microfluidic devices as they form biofilms. Unexpectedly, these experiments reveal that individual bacteria use pili-based (‘twitching’) motility to actively navigate towards regions with higher antibiotic concentrations. Our analyses suggest that this biased migration is driven, at least in part, by a direct response to the antibiotics. Migrating cells can reach antibiotic concentrations hundreds of times higher than their minimum inhibitory concentration in a few hours and remain highly motile. However, isolating these cells - using fluid-walled microfluidic devices that can be reconfigured in situ - suggests that these bacteria are terminal and not able to reproduce. In spite of moving towards their death, we show that migrating cells are capable of entering a suicidal program to release bacteriocins that are used to kill other bacteria. Our work suggests that bacteria respond to antibiotics as if they come from a competing colony growing in the neighbourhood, inducing them to invade and attack. As a result, clinical antibiotics have the potential to serve as a bait that lures bacteria to their death.

## Introduction

Bacterial biofilms often experience steep antibiotic gradients when they are treated because the high cell density within these communities both attenuates diffusion and removes compounds from circulation (1–3). In addition, bacteria commonly live alongside other strains and species that themselves release antimicrobials that diffuse and can again generate chemical gradients (4–6). In order to cope with life under these conditions, bacteria have evolved a wide variety of physiological responses that detect and respond to toxic compounds. It has been hypothesised that many of these responses evolved as a way to cope with competition from other strains and species (‘competition sensing’, (7, 8)). For example, bacteria are known to upregulate production of bacteriocins and antibiotics to generate reciprocal attacks against other toxin-producing strains (9–12). In addition, there are responses that can defend bacteria from toxic compounds, such as the upregulation of efflux pumps and biofilm formation (8, 13), which also serve to protect bacteria from clinical antibiotics (14).

One of the key ways that bacteria physiologically respond to chemicals in their environment is via chemotaxis; the ability of cells to bias their motility in response to chemical gradients (15, 16). There is a large literature on swimming (flagella-driven) chemotaxis, which broadly suggests that cells have the ability to move towards beneficial conditions and away from harmful ones (17–22). However, to our knowledge, there is no evidence that swimming cells bias their movement in response to antibiotics. Much less is known about the motility responses to chemical gradients of surface-attached bacteria, but the opportunistic pathogen *Pseudomonas aeruginosa* is able to bias its twitching motility across surfaces towards nutrients (23). Twitching motility is driven by grappling-hook like pili that extend and retract to pull bacteria over surfaces and other cells. As such, this form of motility is prevalent within dense communities like biofilms where chemical gradients, including antibiotic gradients, are expected to be most pronounced (1–3).

We hypothesized, therefore, that twitching motility would be important for the ability of cells to survive antibiotics within surface-attached communities like biofilms, which are well known for their ability to tolerate antibiotic treatment (24). Specifically, we predicted that bacteria would have evolved to move away from antibiotics, thereby enabling a larger population to survive. Our prediction was wrong: not only do cells not move away from antibiotics, we found that they actively move towards them, killing themselves in the process.

## Results and Discussion

### Surface-attached bacteria move towards antibiotics via twitching motility

We used microfluidic devices and automated cell tracking to quantify the movement of *P. aeruginosa* cells as they are exposed to well-defined spatial gradients of antibiotics in developing biofilms (Fig. 1). We began with the antibiotic ciprofloxacin, which is widely used to treat *P. aeruginosa* infections (25, 26). To set a baseline, we first determined the minimum inhibitory concentration (hereafter MIC) of ciprofloxacin for *P. aeruginosa* (strain PAO1) in shaking cultures, which agrees with the published MIC of this strain (Fig. S1, (27)). We then exposed surface-attached cells to an antibiotic gradient in a microfluidic device where the antibiotic concentration ranged from zero to 10 times the MIC (Fig. 1*A* and *B*, Materials and Methods). After approximately 5 h of unbiased movement, we were surprised to see that twitching cells began to bias their movement towards increasing concentrations of ciprofloxacin (Fig. 1>*B* and *D*, Movie 1). The movement bias, *β*, defined as the number of cells moving up the gradient divided by the number of cells moving down the gradient, peaks after approximately 10 h and then decays as the surface becomes crowded with cells (Movie 1) and tracking becomes difficult (Materials and Methods). We explored gradients that varied in steepness, always starting at a ciprofloxacin concentration of zero and up to a maximum of 0.1, 1, 10, 100 and 1000 times the MIC, whilst keeping the length scales of the gradient constant. Whenever the concentration was higher than the MIC, cells biased their movement towards the antibiotics (peaking at *β* ≈ 2.5 to 4, Fig. 1*D*). These experiments revealed that cells are capable of entering regions of extremely high concentration (up to 1000 times MIC) and remain motile for the remainder of the experiments (≈ 5 h; Fig. 1*D*, S2, Materials and Methods). In contrast, the direction of cell movement in a control without antibiotic was approximately random (*β* ≈ 1, Fig. 1*C*).

**Figure 1.**
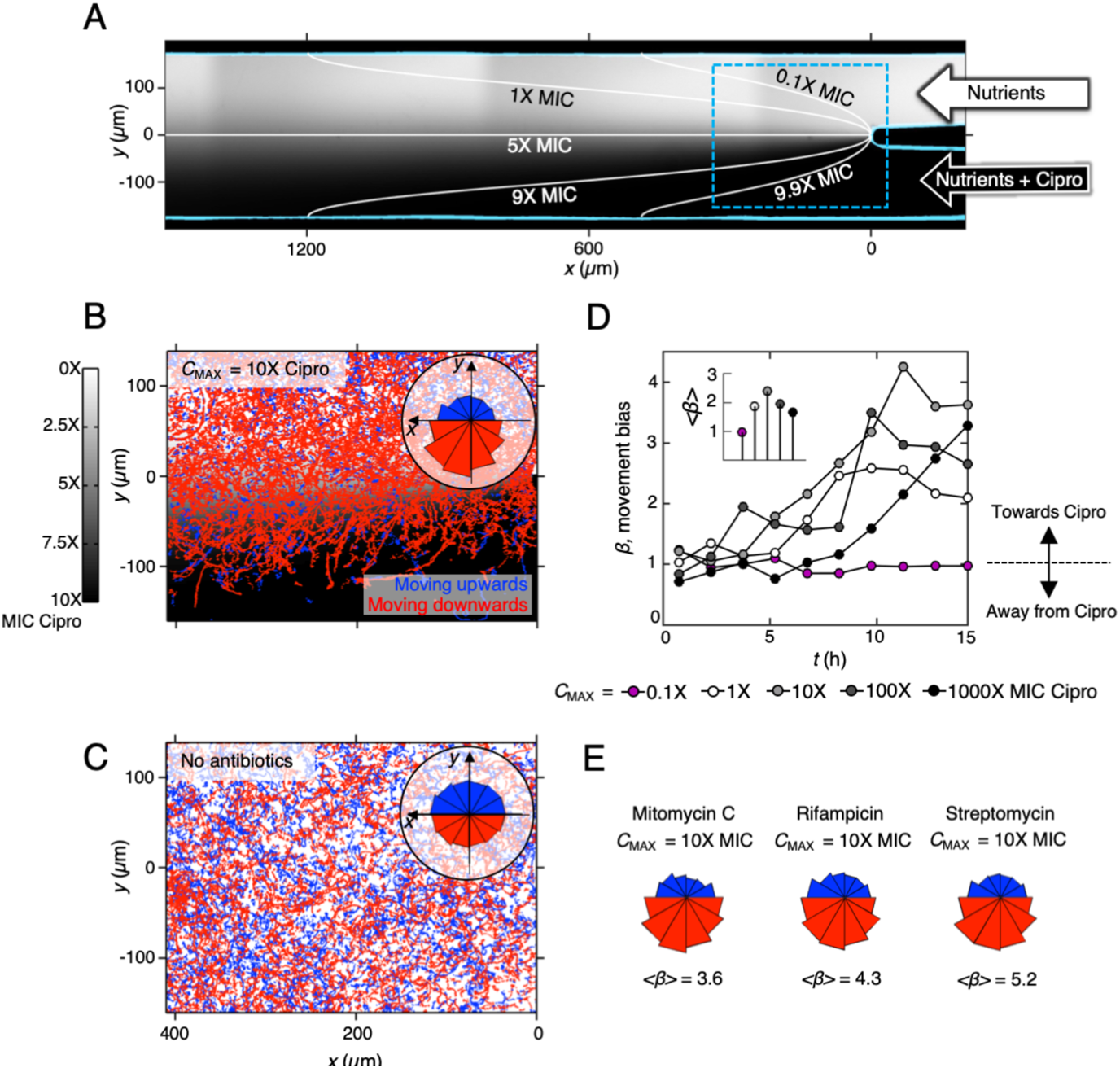
Twitching *P. aeruginosa* cells bias their motility towards increasing concentrations of antibiotics. **(A)** A dual-inlet microfluidic device generates a steady antibiotic gradient via molecular diffusion. Lines show isocontours of ciprofloxacin concentration within the device when *C*_MAX_ = 10X MIC ciprofloxacin is injected through the bottom inlet. The background shading shows approximate distribution of ciprofloxacin, experimentally imaged here using fluorescein, whilst isocontours were calculated using a mathematical model (Materials and Methods). **(B)** Cell trajectories are colour-coded according to whether they move towards increasing (red) or decreasing (blue) ciprofloxacin concentration. Inset: Cell movement direction plotted as a circular histogram reveals that cells bias their motility towards increasing ciprofloxacin concentrations. **(C)** No such bias is seen when nutrient media is added to both inlets so that no antibiotic gradient is present. **(D)** The cell movement bias, *β,* is defined as the number of cells moving up the gradient divided by the number of cells moving down the gradient. The movement of cells is not biased in either direction (*β* ≈ 1) for the first ≈5 h of the experiment, after which cells begin to bias their movement towards ciprofloxacin (*β* > 1). Cells move towards ciprofloxacin even when the ciprofloxacin concentration on one side of the device is 1000 times the MIC (*C*_MAX_ = 1000X MIC, black circles), while biased movement is not present when ciprofloxacin is injected into the device at sub-inhibitory levels (*C*_MAX_ = 0.1X MIC, magenta circles). Inset: The overall bias calculated from trajectories at all time points, <*β*>, shows that *C*_MAX_ = 10X MIC induces the largest response. **(E)** Circular histograms and measurements of <*β*> reveal that cells also bias their movement towards other antibiotics.

This motility response is not limited to ciprofloxacin: we found that cells also bias their movement up gradients of antibiotics belonging to different antibiotic classes that have different chemistries and mechanisms of action (Fig. 1*E*). Ciprofloxacin causes DNA damage by inhibiting an enzyme (gyrase) involved in DNA coiling, but we found that cells also move towards: rifampicin, which targets an RNA polymerase; streptomycin, which targets the ribosome; and mitomycin C, which is again a DNA damaging agent but one that acts by cross-linking DNA. We also tested carbenicillin and other β-lactam antibiotics, but these caused extreme cell elongation that inhibited cell movement in our assay and so we did not study them further (Fig. S3). The diversity in both the chemistry and mechanism of action of the antibiotics that elicit biased movement suggests a general response to the cell damage caused by antibiotics, rather than to the specific antibiotics themselves. Further consistent with the importance of cell damage, we found that sub-MIC concentrations of ciprofloxacin do not bias cell motility (Fig 1D).

The movement of cells in antibiotic gradients appeared to conform to the definition of chemotaxis: the ability of cells to actively bias their motion in response to chemical gradients (16). How, though, are cells biasing their motility towards antibiotics? From previous work, we know that cell movement in our assays is driven by twitching motility and that these twitching cells can actively bias their movement towards beneficial nutrients by reversing their movement direction more frequently when moving away from a nutrient source (23). Is biased motility in antibiotic gradients driven by the same behaviour? To test this hypothesis, we began by quantifying cellular reversal rates and found that cells moving away from ciprofloxacin actively reversed direction more frequently than cells moving towards them (Fig. S4). A previous study showed that cells lacking the key response regulator of twitching chemotaxis, PilG, remain motile but have significantly reduced reversal rates compared to wild-type (WT) cells and are unable to chemotax towards beneficial nutrients (23). Consistent with this previous report, we find here that cells with an in-frame deletion of *pilG* (*ΔpilG*) are also incapable of biasing their movement towards increasing ciprofloxacin concentrations, despite remaining motile in this assay and having an identical MIC as WT cells (Fig. S1, S5 and Movie 2). It is worth noting that the *ΔpilG* cells do show reduced motility in the assay relative to the WT cells but, importantly, our measurements of movement bias only use cells that demonstrate appreciable movement, thus preventing non-motile cells of either genotype from influencing our measurements (Materials and Methods). When this *ΔpilG* strain is complemented at the *pilG* locus, its ability to move towards ciprofloxacin is restored (Fig. S5 and Movie 2). Taken together, these results suggest a common behavioural basis for twitching chemotaxis towards antibiotics and nutrients.

### Nutrient gradients do not explain chemotaxis towards antibiotics

Cells only begin to bias their motility up antibiotic gradients after an initial ≈5 h period of nearly random motility (Fig. 1*D*). This delayed response introduces an important complication, as secondary gradients in nutrients and other compounds potentially released by cells are expected to build up in the device over time, which might indirectly drive movement towards antibiotics. In particular, cells situated in regions of the device with lower antibiotic concentrations can rapidly proliferate, while cells initially in regions with higher antibiotic concentrations either tend to detach or die (Movie 1). The different numbers of cells on either side of the device, both within the test section (Fig. 1*A*) and in the regions upstream, means that emergent chemical gradients can form because the cells consume nutrients and release compounds at different rates on either side of the device. Thus, cell movement towards higher antibiotic concentrations could be, in principle, driven by movement towards higher nutrient concentrations and towards lower concentrations of the diverse set of compounds released by cells. We therefore sought to confirm whether such secondary gradients could be responsible for the biased movement we observe, rather than it being a direct response to the antibiotic gradients themselves. We first tried switching the direction of the antibiotic gradient, after the cells had established themselves in the regions with lower antibiotic concentrations. However, the sudden change in antibiotic concentration caused cells to either stop moving or detach altogether. We therefore sought alternative approaches to control for the formation of secondary *de novo* gradients.

We focused first on whether putative emergent nutrient gradients could be responsible for the observed biased movement towards antibiotics. Previous work has shown that surface-attached *P. aeruginosa* cells undergo chemotaxis up gradients of the metabolizable carbon source succinate (23). Consistent with this, we found that twitching cells will also bias their motility up gradients of tryptone, the growth media used in this study (Fig. 2*A*), a response that has also been observed in swimming *P. aeruginosa* cells (28). To test whether the biased movement we see in our experiments with antibiotics could be driven solely by *de novo* gradients in tryptone, we created opposing gradients of tryptone and ciprofloxacin in our microfluidic device. Specifically, we injected full strength tryptone through one inlet of the device and ciprofloxacin mixed with tryptone at 10% of the regular concentration through the other inlet. We use 10% media rather than 0% because with the latter, we find that cells quickly stop moving in regions without nutrients. If cells in our previous experiments are simply moving in response to nutrient gradients, we expect that cells would now move away from the antibiotic source. While cells initially move in the direction of increased tryptone, after ≈5 h this response rapidly drops off and after ≈7.5 h cells again exhibit biased movement in the opposite direction towards increasing antibiotic concentrations (Fig. 2*A*). The robust movement of cells towards low nutrient and high antibiotic concentrations so early in the experiment suggest that chemotaxis towards antibiotics (Fig. 1) is not driven by nutrient gradients.

**Figure 2.**
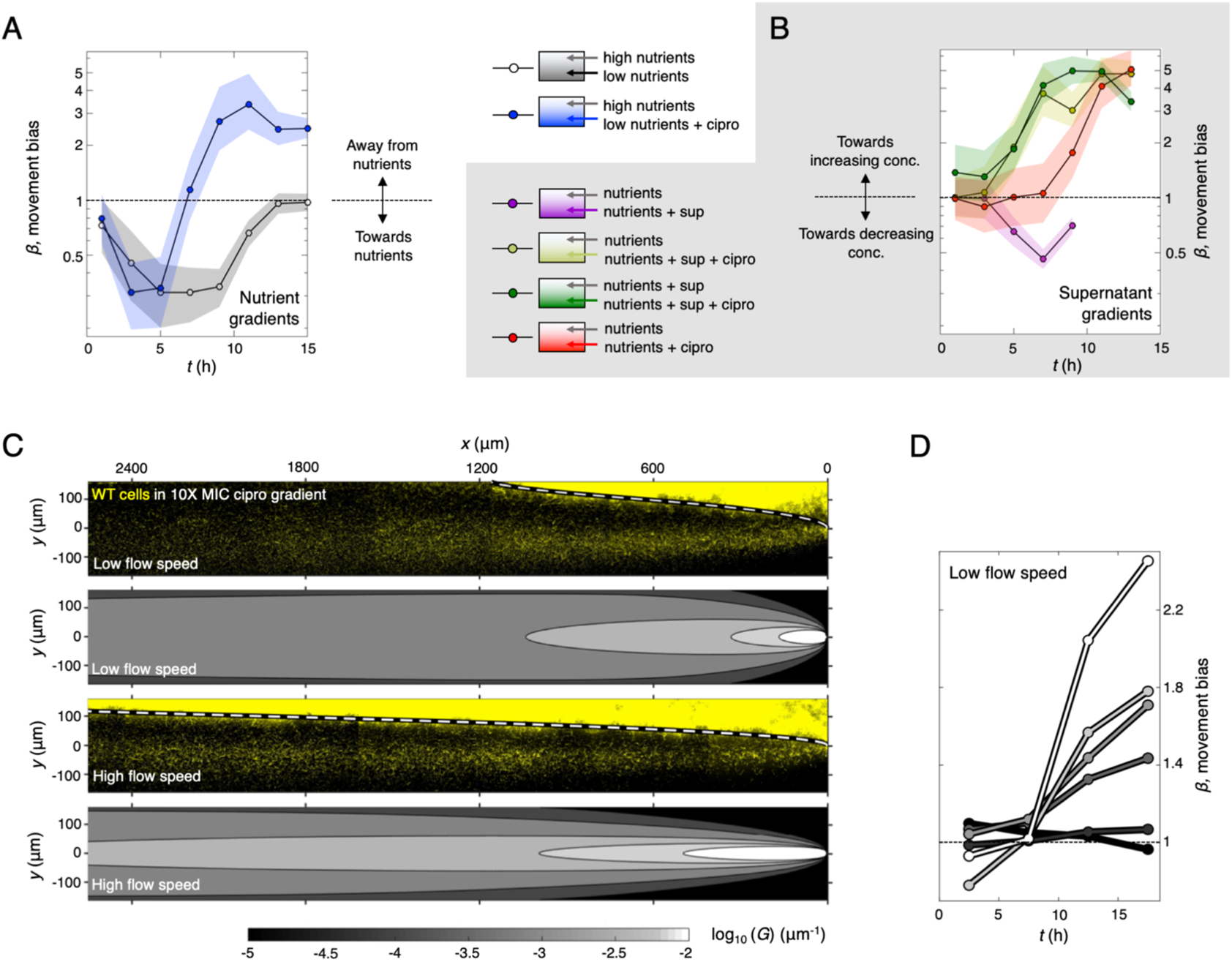
Putative gradients of nutrients or cell products do not explain movement towards antibiotics. **(A)** When exposed to a gradient of tryptone (*C*_MIN_ = 10% to *C*_MAX_ = 100% of the regular concentration used in our growth medium), cells bias their motility towards increasing tryptone concentrations (black line). However, when we add ciprofloxacin (*C*_MAX_ = 10X MIC) to the side of the device that contains the lower concentration of tryptone, we observe cells initially bias their movement towards increasing tryptone concentrations for the first ≈7.5 h, after which cells begin to bias their motility in the opposite direction so that they move towards increasing ciprofloxacin concentrations (blue line). **(B)** When exposed to a gradient of cell-free supernatant (*C*_MAX_ = 10%), cells bias their motility away from the supernatant (purple line). However, when ciprofloxacin (*C*_MAX_ = 10X MIC) is added to the supernatant, cells rapidly (after ≈5 h) bias their motility towards increasing ciprofloxacin concentrations, even though this now results in them moving towards the supernatant (light green line). Furthermore, if cells are exposed to a gradient of ciprofloxacin with a uniform background concentration of 10% cell-free supernatant, they also undergo chemotaxis towards ciprofloxacin (dark-green). Indeed, in the presence of supernatant, cells bias their motility towards ciprofloxacin even more rapidly than in the absence of supernatant. **(C)** After ≈20 h, YFP-labelled WT cells growing in a gradient of ciprofloxacin (*C*_MAX_ = 10X MIC) form a dense biofilm wherever the local concentration of antibiotic is <1X MIC, whilst a smaller band of migrating cells can be observed in regions where the local concentration of antibiotic is much larger. Increasing the flow through the device by threefold sharpens the strength of the antibiotic gradients, allowing both the 1X MIC isocontour (dashed white line) and band of migrating cells to stretch further downstream. Here grey regions show the magnitude of the normalised antibiotic gradient, *G =* 1/ *C*_MAX_ ∂*C/*∂*y.* **(D)** Measurements of bias for each of the greyscale regions shown in *C* (bottom panel) show that the strength of chemotaxis peaks where the antibiotic gradient is the largest. Here, the white line corresponds to the bias of cells in the region of the device where the gradient is strongest, whereas the black line indicates the bias of cells in region where the gradient is weakest. This analysis used data from the higher flow rate experiment because it allowed us to collect data from a larger area of the device.

### Repulsion by cell-free supernatant does not explain chemotaxis towards antibiotics

We next explored the possibility that biased movement in antibiotic gradients could be explained by products released by cells that might trigger repulsion away from regions of the device with high cell density. Consistent with the potential for repulsion from areas of high cell density, we find that twitching cells are repelled by cell-free supernatant extracted from cells grown to high density under static conditions in the multi-well plate assays that are widely used to study biofilm physiology (Fig. 2*B*, Materials and Methods (13, 29)). Specifically, we mixed cell-free supernatant collected from static well-plates with fresh media (20% supernatant, 80% tryptone broth) and injected it through one inlet of our device, whilst media mixed with the same proportion of water (20% water, 80% tryptone broth) was injected through the other inlet. These experiments showed that cells move away from the channel containing supernatant (Fig. 2*B*, purple line). This result is interesting as it suggests that in *P. aeruginosa*, twitching motility will guide cells away from regions of high cell density in developing biofilms, potentially facilitating the colonization of new territory. However, it also raises the possibility that this repulsion could explain the migration towards antibiotics. We therefore performed a number of additional experiments to explore this possibility.

We reasoned that if a factor released by cells is responsible for movement towards antibiotics, then adding cell-free supernatant in the background of an antibiotic gradient should limit or stop the movement of cells toward higher antibiotic concentrations. We tested this hypothesis with two additional experiments. The first experiment exposed cells to a gradient of ciprofloxacin in a uniform background of cell-free supernatant. We again used cell-free supernatant collected from high-density static cultures at 20% concentration, which we expected to saturate any gradients in cell products that might occur *de novo* in the microfluidic device where cell densities are initially low (Fig. 2*B*, green line). The second experiment exposed cells to a ciprofloxacin gradient and supernatant gradient simultaneously, with the larger concentration of both on the same side of the device, so that repulsion of cells away from supernatant would oppose the putative movement towards ciprofloxacin (Fig. 2*B*, yellow line). We still observed movement towards ciprofloxacin in both of these experiments. In fact, we found that movement towards ciprofloxacin actually commences more rapidly in the presence of the cell-free supernatant than without (Fig. 2*B*, red line). Importantly, this rapid migration towards antibiotics occurred even when the cell-free supernatant gradient was oriented such that one would expect it to repel cells away from the antibiotics. In sum, repulsion due to factors present in cell-free supernatant could promote cell movement from regions of high cell density towards antibiotics, but this effect does not explain the observed chemotaxis towards ciprofloxacin.

There remains the possibility that chemotaxis towards antibiotics is driven by a repulsive factor, or set of factors, that are not present in the cell-free supernatant used in the above experiments, which was collected from static cultures grown in multi-well plates. For example, it may be that cells growing under flow in the microfluidic device have a different physiology, such that any secondary chemical gradients that develop within the device are not captured by adding cell-free supernatant collected from static cultures on the bench. We therefore repeated the cell-free supernatant experiments using media collected directly from cells growing within microfluidic devices (Fig. S6, Movie 3, Materials and Methods). To maximise the likelihood of observing a response, we used 100% cell-free supernatant for these experiments and flowed this through one inlet with growth media (tryptone broth) through the other. This revealed that, as for the standing culture experiments, cells move away from the channel with cell-free supernatant obtained from microfluidic devices. With 100% cell free supernatant, this response is likely to be a combination of both moving away from cell-free supernatant and towards an increasing nutrient concentration. When we combined the cell-free supernatant with 10X MIC ciprofloxacin, cell movement was initially biased away from the cell-free supernatant/ciprofloxacin. However, importantly, after ≈7 h, cells began to move in the opposite direction, towards the cell-free supernatant and ciprofloxacin. The addition of cell-free supernatant on the antibiotic side of the device, therefore, does not prevent chemotaxis towards the antibiotic, as would be expected if cell products released in the device were driving the response. These experiments again suggest that repulsion by cell free supernatant is not sufficient to explain migration towards antibiotics.

### A factor released in response to antibiotics does not explain chemotaxis

There remains the possibility that chemotaxis is driven by a repulsive factor that is not present in the cell-free supernatants we studied. These supernatants were collected from experiments without antibiotic present, as any residual antibiotic remaining in the collected supernatant would complicate interpretation. However, this leaves open the possibility that surface-attached cells within our microfluidic experiments might release a compound in response to the antibiotics (i.e. antibiotic-induced cell products) that could drive directional cell movement, something that has been observed for swarming bacteria (30). To examine this possibility, we used a classical result from the chemotaxis literature: movement bias is predicted to increase with the strength of the chemical gradient. This correlation has not only been demonstrated in chemotaxis occurring in twitching bacteria (23), but also in swimming bacteria (15) and eukaryotic cells (31). Since the distribution of ciprofloxacin in our microfluidic devices is predicted to differ starkly from that of putative antibiotic-induced cell products (Fig. S7 and S8), measuring how movement bias changes at different positions within the device gives us the ability to distinguish which of these alternatives is most likely to explain the observed patterns of cell movement. A model of diffusion was used to quantify how the concentration of ciprofloxacin, and its spatial gradient, varies within our device (Materials and Methods). The isocontour corresponding to the MIC of our strain (1X MIC) was found to closely match the position where biofilm growth becomes strongly suppressed by antibiotics (dashed line, Fig. 2*C*), which is a benchmark that supports the accuracy of our model. If the flow rate within the device is increased by a factor of three, the 1X MIC isocontour was predicted to be pushed further downstream, which again closely matches the distribution of biofilm observed experimentally under these higher flow conditions (Fig. 2*C*).

Next, we calculated the movement bias, *β,* in five different regions of our device, each corresponding to different strengths of ciprofloxacin gradient. These analyses used higher flow rate experiments because the gradients are sharpened, allowing us to simultaneously image cells in four different fields of view to collect more cell trajectories within each gradient strength bin (Fig. 2*C*, Materials and Methods). Our analyses reveal that the strength of the ciprofloxacin gradient (*G* = 1/*C*_MAX_ ∂*C/*∂*y*) is an excellent predictor of *β* - after ten hours, *β* was found to monotonically increase with *G* (Fig. 2*D*). These results indicate that the stimulus the cells are responding to closely matches the predicted gradient of ciprofloxacin within our device. While the gradient of ciprofloxacin decreases as one moves downstream, the gradient of a putative antibiotic-induced cell product would likely increase in the downstream direction. This is because as fluid passes through the device, it passes by more and more cells exposed to sub-MIC levels of antibiotic, which would steadily increase the concentration of the product on one side of the device (Fig. S7 and S8). While the exact distribution of a putative cell product is hard to predict, the fact that it would differ strongly from that of ciprofloxacin suggests that the excellent correlation between *G* and *β* (Fig. 2*D*) is unlikely to be driven by antibiotic-induced cell products.

Taken together, our data suggests that neither nutrient depletion, nor products released either in the presence or absence of antibiotics, can explain the movement of cells towards ciprofloxacin. These results suggest that we are observing a direct response to the antibiotics. There is a growing body of work showing that flagella-based swimming can transport bacteria into regions of high antibiotic concentration (32–35). However, to the best of our knowledge, there is no evidence of chemotaxis in response to the antibiotic gradients themselves. Here, we observe that cells move towards antibiotics, actively reverse direction to facilitate this directed movement and, finally, that movement bias increases monotonically with the strength of the antibiotic gradient, which are all signatures of chemotaxis (15, 16).

### Cells migrate towards antibiotics and die

Our data show that *P. aeruginosa* cells will bias their motility to move into regions with extremely high antibiotic concentrations. Moreover, many cells remain motile at concentrations many times greater than that needed to prevent their growth in a standard liquid culture assay (Fig S2). We therefore wanted to establish the fate of these cells: do they remain viable and gain tolerance to high concentrations of antibiotics, or are they terminal and doomed to die? Or indeed, is it possible that some cells evolve mutations that confer resistance in the short time-scale (≈20 h) of our experiments (32, 34)? The challenge with answering this question is that the cells in question are fully encased within PDMS-based microfluidic devices, which means it is not possible to isolate migrating cells and study them further. One can watch them for the duration of the experiment, but after ≈20 h, the devices become clogged with growing cells. We are therefore unable to ascertain whether cells that migrate into regions of high antibiotic concentration remain active and retain long-term viability.

In order to overcome this challenge, we moved to a novel assay that uses open “fluid-walled” microfluidics (36), where twitching cells can be studied in a chemical gradient and then isolated after they have migrated, (Materials and Methods; for a detailed description of our approach, see (37)). Briefly, microfluidic devices were inoculated with a 1:1 co-culture of YFP-labelled WT cells and unlabelled, chemotaxis-null *ΔpilG* cells, which allowed us to follow cells that can and cannot perform chemotaxis. We then followed cell movement under the microscope until a substantial number of WT cells had begun to migrate towards ciprofloxacin (*C*_MAX_ = 100X MIC, *t* = 24 h; Fig. 3*A*-*C*). To reconfigure the fluid-walled channels, we mounted a PTFE (polytetrafluoroethylene) stylus (38) to the microscope’s condenser and moved the device relative to the stylus using the microscope’s motorised stage. This technique allowed us to reconfigure the central channel into segments in real time and with a high level of accuracy (tens of micrometers). Thus, we could physically isolate the cells that had performed chemotaxis within a fluid-walled chamber (Fig. 3*D* and *E*). Importantly, these cells were strongly biased towards WT rather than the chemotaxis-null *ΔpilG* cells (mean = 84% WT, with 95% confidence interval of 81-87%), confirming our ability to isolate cells that could undergo chemotaxis. To analyse the viability of these isolated cells, we extracted the contents of the chambers and plated them onto agar plates without antibiotics to monitor whether any of the cells were able to form viable colonies. Additionally, we replaced the chamber contents with fresh growth media without antibiotics to monitor the health of any cells that remained attached to the surface of the microfluidic chambers.

**Figure 3.**
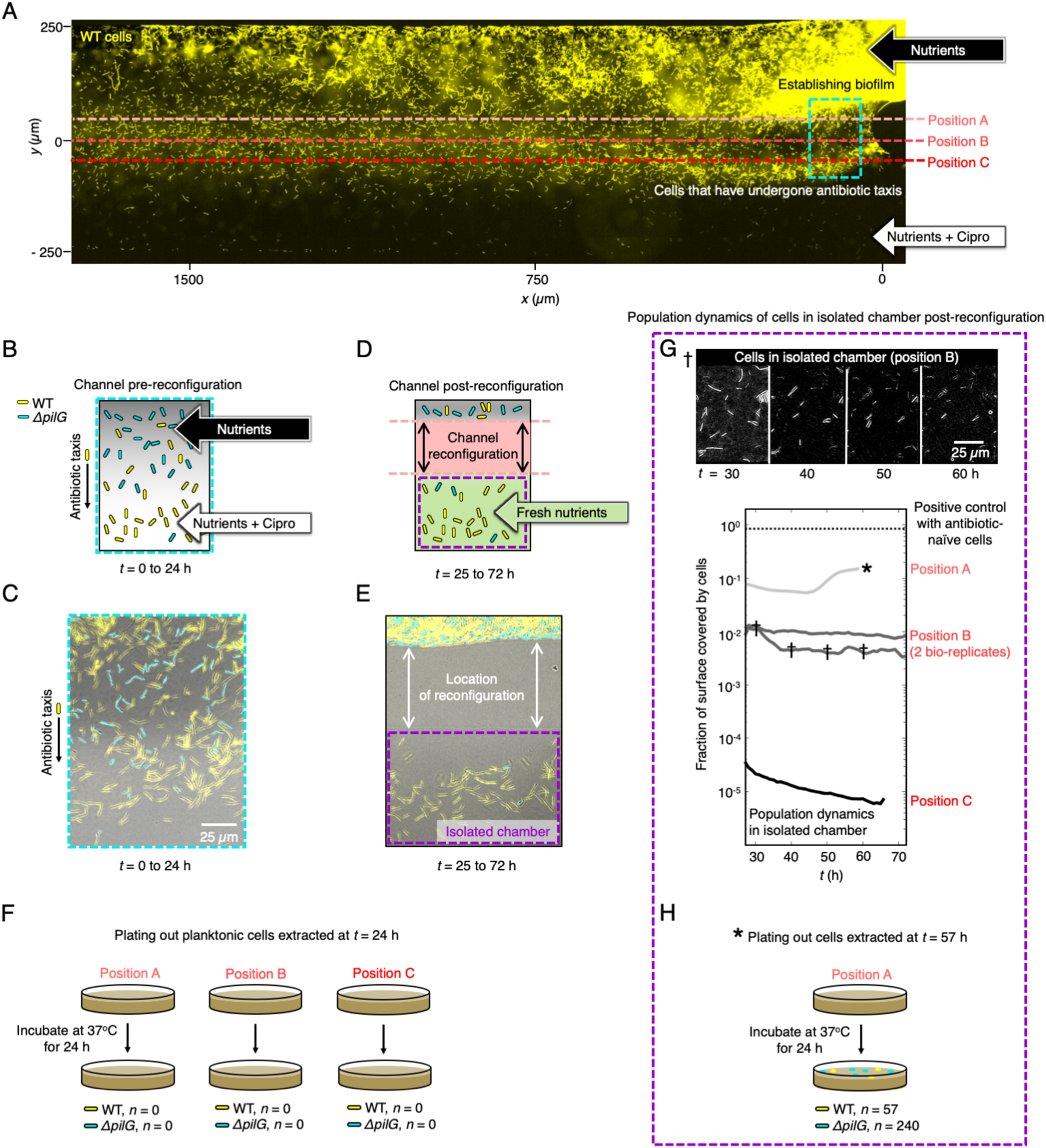
Cells migrate towards antibiotics and die. **(A)** Chemotaxis in a fluid-walled microfluidic device, which can be reconfigured in real-time, allowing us to isolate and extract the chemotaxing cells. Equal numbers of YFP-labelled WT cells (yellow) and unlabelled, chemotaxis-null mutant cells (*ΔpilG,* not-shown in this YFP image) were inoculated and exposed to a gradient of ciprofloxacin (*C*_MAX_ = 100X MIC). To test cell viability in different parts of the device after *t* ≈ 24 h of incubation, we reconfigured the channel at three different locations, shown with the dashed lines, across separate experiments. ‘Position A’ isolated all chemotaxing cells, including those that had only just begun moving towards ciprofloxacin. ‘Position B’ was more conservative in that cells had to chemotaxis further to be captured. ‘Position C’ was more conservative still and isolated only those chemotaxing cells that had reached ciprofloxacin concentrations close to *C*_MAX_. **(B)** Cartoon of channel prior to reconfiguring the channel: we predicted that WT cells (shown in yellow) would move towards higher ciprofloxacin concentrations (lighter shades of grey), whilst *ΔpilG* cells (shown in cyan) would be unable to bias their motility in this manner. **(C)** The microfluidic device imaged at the approximate location of the dashed cyan box in (A) immediately prior to reconfiguring the channels (*t* ≈ 24 h) confirmed our prediction. **(D)** Cartoon of channel after reconfiguring the channel with PTFE stylus: the reconfiguration was expected to create a microfluidic chamber containing mostly WT cells and fresh nutrient media (shown in green) was used to replace any residual antibiotics remaining in this chamber. **(E)** Experimental image of the same region of the microfluidic device shown in (C) after it was reconfigured shows that, as expected, the isolated fluid chambers contained mostly WT cells (mean = 84%, with a 95% confidence interval of 81-87%). **(F)** Planktonic cells were extracted from the chamber, plated onto nutrient agar, and monitored for growth. No cell growth was detected on these plates in any of our experiments, indicating that these planktonic cells were not viable. **(G)** However, many cells remained in the isolated chambers attached to the surface, which we subsequently imaged for a further ≈40 h. In experiments where the channel was reconfigured at Position B (dark-grey lines, two biological repeats) or C (black line), no cell growth was detectable and indeed the fraction of the surface covered by cells decreased steadily as cells continued to die or detach from the surface. This can also be seen in the representative sequence of images (reconfiguration made at Position B, marked by †), illustrating how the number of cells attached to the surface decreases over time due to cell death and detachment. When the reconfiguration was made at Position A, the fraction of the surface covered by cells initially decreased, momentarily increased, and finally plateaued at approximately 10% coverage. After *t* ≈ 60 h (see *), we extracted the entire contents of the well and plated it onto nutrient agar plates. **(H)** After a further 24 h, we recovered a small number of both WT and *ΔpilG* colonies, which suggested that the reconfiguration made at Position A had isolated some cells that were not undergoing chemotaxis and had only been exposed to relatively low ciprofloxacin concentrations.

We reasoned that cell viability might depend on how far a cell has migrated up the ciprofloxacin gradient and thus the ciprofloxacin concentration it has experienced. Therefore, we carried out three separate experiments in which the microfluidic channels were reconfigured at different positions, allowing us to isolate three different sub-populations of cells that had each experienced a different minimum concentration of ciprofloxacin. In the first position, we reconfigured the channel to isolate only those cells that had already undergone chemotaxis and thus begun to accumulate in regions of high ciprofloxacin concentration (“Position C”, Fig. 3*A*). The cells extracted from this sub-population were found to be incapable of forming colonies on agar (Fig. 3*F*) and none of the cells that remained within the isolated fluid chamber underwent any further cell division in over 36 h (Fig. 3*G*). To confirm that the fluid chamber provided a suitable environment for cell growth, we inoculated ≈25 healthy cells into the chamber at the end of the experiment. These freshly-inoculated cells grew rapidly, completely covering the bottom surface of the chamber within 24 h (dotted line, Fig. 3*G*), demonstrating that the chamber did not contain residual antibiotics that could stifle growth. In the second position, we reconfigured the channel to isolate both cells that had already undergone chemotaxis as well as the majority of cells that were still migrating (“Position B”, Fig. 3*A*). Once again, none of these cells showed evidence of long-term viability (Fig. 3*F, G*). Finally, we reconfigured the channel to isolate all cells that were undergoing or had already undergone chemotaxis, including those that had only very recently begun to start moving up the ciprofloxacin gradient (“Position A”, Fig. 3*A*). Once again, the extracted cells were found to be incapable of forming colonies (Fig. 3*F*). Whilst there was initially no detectable cell growth within the fluid chamber, after approximately 20 h, the early stages of a biofilm containing both WT and *ΔpilG* cells was observed forming on the surfaces of the fluid chamber (Fig. 3*G*). After a further 10 h, the growth of this biofilm appeared to have stalled and so the entire chamber contents were extracted and plated onto antibiotic-free nutrient-rich agar plates, on which a relatively small number of colonies were recovered (Fig. 3*H*). Interestingly, while the majority of cells seen to migrate towards antibiotics were WT, the majority of colonies that grew were chemotaxis-null *ΔpilG* cells, which were less likely to enter the regions of high antibiotic concentration. Taken together, these three separate channel reconfigurations suggest that whilst some cells retain long-term viability at the earliest stages of chemotaxis, they will continue to move into higher antibiotic concentrations where they rapidly lose this viability and will ultimately die.

### Twitching bacteria move towards supernatant from competitors

Why does *P. aeruginosa* perform an ultimately fatal migration towards antibiotics? We hypothesized that the observed migration towards clinical antibiotics may have its evolutionary origins in the natural ecology of *P. aeruginosa*. Clinical antibiotics are widely reported to induce biofilm formation and this can be recapitulated in *P. aeruginosa* using the cell-free supernatant of other *P. aeruginosa* strains (13). However, this effect is only observed when the supernatant is toxic to the focal strain, which is consistent with the hypothesis of competition sensing: an evolved ability to detect and respond to harmful competing strains (7). Under this hypothesis, the presence of clinical antibiotics causes bacteria to act as though the attack is coming from ecological competitors (13).

We therefore asked whether, in a similar way to their response to clinical antibiotics, cells might bias their movement up gradients of toxic cell-free supernatant isolated from different *P. aeruginosa* strains. Based on previous work, we selected one strain (‘Strain 7’) that strongly inhibits the growth of our focal strain PAO1 in coculture and a second strain (PA14) that does not (Fig. 4*A*, (13)). Consistent with our hypothesis, we observed biased movement towards the toxic supernatant that stifles cell growth, but unbiased movement in a gradient of supernatant from the non-toxic strain (both using a supernatant concentration of 10%, Fig. 4*A*). These data again suggest that the response we observe with clinical antibiotics is a general response to a gradient of toxic components. But why would cells move towards toxic compounds? Moving towards toxic stimuli may be a maladaptive response driven by their adverse effects on cells. Alternatively, it may constitute part of a general strategy to attack and invade the territory of neighbouring genotypes that present a threat. Consistent with the latter, the evolution of mass suicide to release toxins was recently documented in non-motile *Escherichia coli* (9). Such aggressive strategies also have a precedent in the animal world, particularly in the social insects that are well known for territorial behaviours, where workers will actively seek out and attack neighbouring colonies and other harmful species, and die in the process (39).

**Figure 4.**
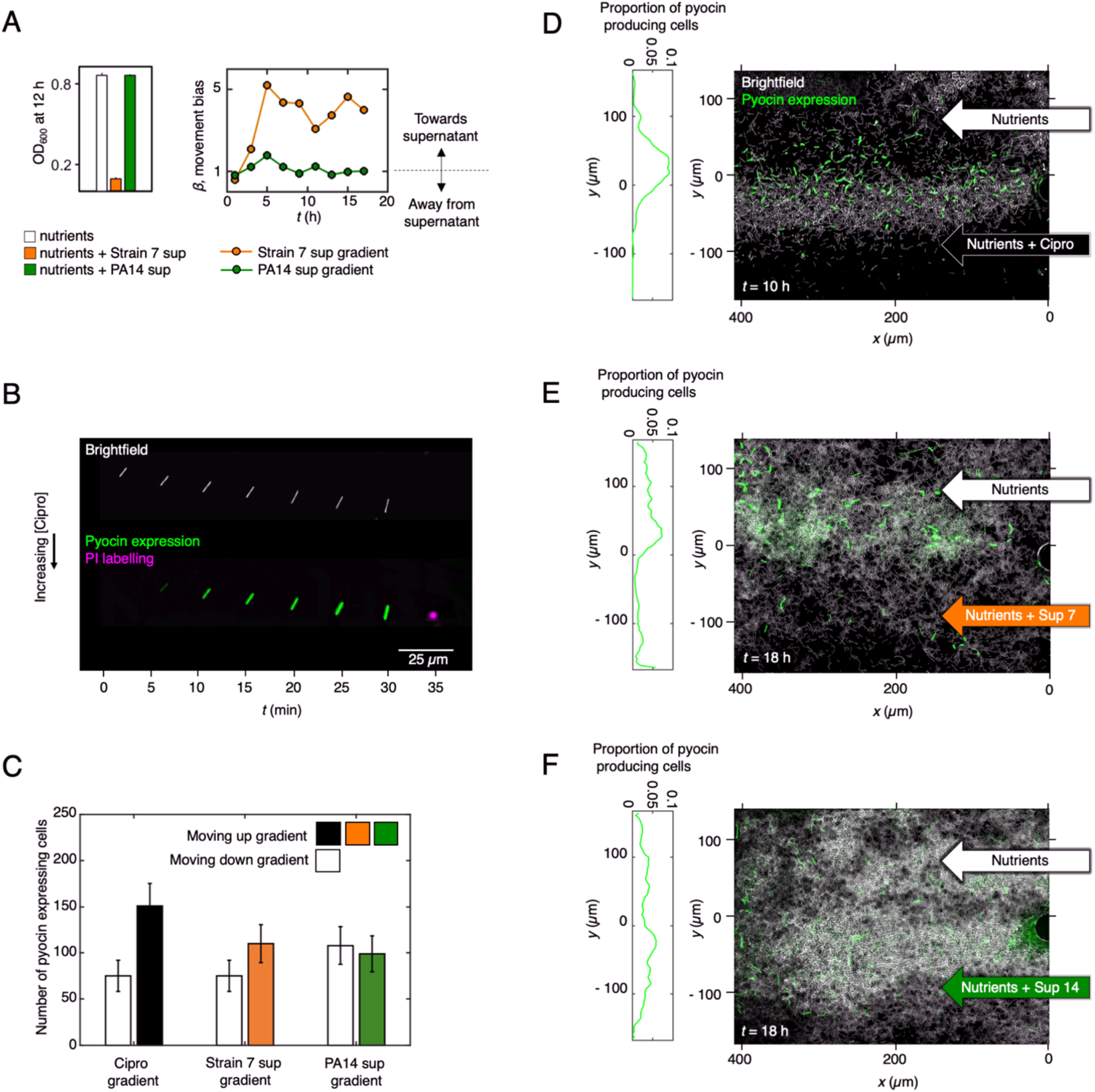
Cells produce bacteriocins (pyocins) as they move towards both antibiotics and toxic supernatant collected from a competitor strain. **(A)** The growth of our focal PAO1 strain is inhibited by cell-free supernatant extracted from cultures of the soil isolate Strain 7 (orange bar) but is unaffected by that from the clinical isolate PA14 (dark-green bar). When exposed to gradients of the cell-free supernatant of these two different *P. aeruginosa* isolates, our focal PAO1 strain biases its movement towards the toxic supernatant from Strain 7 (*β* > 1, orange line), but displays unbiased movement in gradients of supernatant from the non-inhibitory PA14 (*β* ≈ 1, dark-green line). **(B)** We engineered a reporter strain of PAO1 to allow us to monitor pyocin expression using mNeonGreen fluorescent protein. Here we show a timeseries of a representative cell undergoing chemotaxis towards ciprofloxacin as it gradually turns on pyocin expression (mNeonGreen is shown in green, whilst the corresponding brightfield images have been processed so that cells appear white). The cell eventually undergoes lysis at *t* = 35 min, which can be seen as a transient burst of propidium iodine (shown in purple), which fluorescently labels DNA released from the cell. **(C)** In gradients of ciprofloxacin (*C*_MAX_ = 10X MIC, black bar) and toxic supernatant (Strain 7 Sup, orange bar), a larger number of pyocin expressing cells move towards larger concentrations of the chemoattractant, rather than towards smaller concentrations (*Z*-test for proportions yield *p* < 0.001 and *p* < 0.01 respectively), whilst such a bias is absent in gradients of non-toxic supernatant (PA14, dark-green bar, *p* > 0.7). Counts shown are pooled across two bio-replicates and error bars show 95% confidence intervals. **(D)** After being exposed to a gradient of ciprofloxacin (*C*_MAX_ = 10X MIC) for 10 h, many cells (cell bodies shown in white) have undergone chemotaxis towards increasing ciprofloxacin concentrations and often express pyocins as they do so (quantified using mNeonGreen fusion). The proportion of cells expressing pyocins peaks towards the middle of the microfluidic channel, where the ciprofloxacin gradient is relatively steep. **(E)** Similarly, many of the cells chemotaxing towards toxic supernatant from an inhibitory *P. aeruginosa* Strain 7 (orange arrow) also express pyocins. **(F)** Pyocin expressing cells are also seen in the presence of non-toxic supernatant from a non-inhibitory *P. aeruginosa* strain PA14 (dark-green arrow) but, in contrast to the other conditions, cells do not bias their movement here.

In this model, *P. aeruginosa* has evolved to move towards competitors in order to engage in a counter-attack. If this model is correct, we reasoned one should see the release of antimicrobials by *P. aeruginosa* as it migrates up the gradient of antibiotics or toxic cell-free supernatant. In order to test this hypothesis, we built a fluorescent transcriptional reporter for Pyocin R2 of *P. aeruginosa*, which is induced by DNA damage and released via cell lysis (40). We observed this strain in gradients of ciprofloxacin and either the toxic, or non-toxic, cell-free supernatant of another *P. aeruginosa* strain. These experiments confirmed that many cells expressed pyocins and often performed an explosive lysis as a result (Fig. 4*B-F*, see also (41)). These lysis events can be seen clearly in the presence of propidium iodine which stains the free DNA released upon lysis, observed as a transient puff of colour (Fig. 4*B*, Movie 4). When combined with chemotaxis, the result is the biased movement of pyocin-releasing cells towards the source of toxins (Fig. 4*C*). The patterns of pyocin release are particularly striking in the case of ciprofloxacin (Fig. 4*D*, Movie 4). This observation fits with the known regulation of pyocins, which is DNA damage dependent (27, 40), and suggests that DNA damaging agents (like ciprofloxacin) will lead to a particularly aggressive response. Consistent with this, the release of colicin bacteriocins in *E. coli* is also known to be upregulated by DNA damaging toxins, albeit in stationary cells (9). Altogether, our experiments suggest that *P. aeruginosa* has evolved to bias its twitching motility towards harmful compounds while releasing bacteriocins, which is consistent with a counter-attack behaviour.

## Conclusion

Our experiments have revealed a surprising response to antibiotic gradients in the pathogen *P. aeruginosa*. Rather than moving away from toxins, as we hypothesised would happen, cells actively bias their motion to undergo chemotaxis towards lethal concentrations of antibiotics. We observe this response for a range of different clinical antibiotics and for inhibitory supernatant from foreign *P. aeruginosa* cells. This finding suggests that the biased movement is driven by a general effect of the cell damage caused by antibiotics, rather than a specific response to the compounds themselves. The weight of our evidence also suggests that cells are responding directly to toxin gradients. Until the mechanistic basis of the response is understood, there will remain the possibility that a secondary effect of the antibiotics is driving the cell migration patterns that we observe. However, the effects of chemotaxis are clear either way - in the presence of a spatial gradient cells change their behaviour, move towards toxins, and undergo bacteriocin production. This behaviour is consistent with a robust response of *P. aeruginosa* to the threat of ecological competition from other bacteria. *P. aeruginosa* is already known to be a strong retaliator from its use of the type six secretion system, which it activates multiple times in response to a single incoming attack from a competitor (42, 43). It is also notable as a species that carries a particularly large number of antibacterial weapons (39). A growing body of evidence, therefore, suggests that *P. aeruginosa* is a highly aggressive species that will attempt to eliminate competitive strains and species growing in its neighbourhood. However, in the context of clinical antibiotics, this strategy works against this species, as the drugs simply lure the cells to their death.

## Materials and Methods

### Strains, Media and Culture Conditions

The main strain used throughout this work is a WT *P. aeruginosa* PAO1 from the Kolter collection, ZK2019 (44). The *P. aeruginosa* strains used as competitors (Fig. 4), are also from the Kolter collection and have been published and described elsewhere (13). They include the clinical isolate PA14 (ZK2466) and the environmental isolate ‘Strain 7’ (ZK3098, (13)), which notably is distinct from the strain PA7 used in other studies (45). To study the role of cellular reversals in driving biased movement towards antibiotics (Fig. S4-6), we used strains with an in-frame deletion mutation in *pilG* (*ΔpilG)* and a strain complemented at the *pilG* locus, both of which have been published and described previously (Bertrand, West and Engel, 2010). For co-culture experiments and to confirm that strain background was not an issue (Fig. 3), we also generated an in-frame *pilG* deletion in our (Kolter) PAO1 WT strain background using allelic replacement vector pJB118 published and described previously (46). This strain was used in co-culture experiments with a YFP-labelled variant of our PAO1 WT, which has also been published and described elsewhere (13). To study pyocin expression (Fig. 4), the mNeonGreen (47) gene, with a ribosome binding site (CTAGATTTAAGAAGGAGATATACAT) was inserted immediately after the stop codon of PA0631 (the final gene of the pyocin R2 operon) by two-step allelic exchange. A construct including the mNeonGreen gene flanked by ≈400bp up and downstream of the insertion site was synthesized (IDT) and cloned into the MCS of pEXG2 (48) using NEB Hifi assembly (NEB). This construct was introduced into our (Kolter) PAO1 WT mentioned above by standard methods (49) and confirmed by sequencing using primers flanking the insertion site (GCGGTGCTGTCATGAGCC/ GGATGTCACCTGCAACCTCA).

All bacterial cultures were grown the day before experiments in shaken LB media at 37**°**C. On the day of the experiment, these were sub-cultured to obtain cells in exponential phase. These were then diluted in tryptone broth (TB, 10 g Bacto tryptone per 1 L water), the growth medium used in all of our microfluidic-based experiments. To find the minimum concentration of each antibiotic that inhibits growth in homogeneous cultures, we used the same growth medium but supplemented with different antibiotic concentrations. Exponential phase cells from overnight cultures were inoculated in flat-bottomed 96-well plates (Nunc) in 200 μl of TB at a starting optical density (at 600 nm) of 0.05, and then were placed inside a plate reader (Tecan, Infinite M200 Pro) for 20 h, where they grew at 22°C under shaken conditions. Measurements were recorded every 30 min and the MIC was defined as the minimum concentration of antibiotic that prevents detectable growth after 20 h (Figure S1).

### Microfluidic experiments

All microfluidic experiments were imaged using a Zeiss Axio Observer inverted microscope with a Zeiss 20X Plan Apochromat objective, Zeiss AxioCam MRm camera, a Zeiss HXP 120 light source, a Zeiss Definite Focus system, and Zeiss filter sets (38 HE, 46 HE, and 63HE) except for the experiment shown in Fig. S6, which was imaged using a Nikon Ti-E inverted microscope with a Hamamatsu Flash 4.0 v2 camera, Nikon Perfect Focus system and a Nikon 20X Plan Apochromat objective. We maintained the temperature at 20-22**°**C, but when our microscope suite was outside of this range, we used a custom-designed microscope incubation chamber with both heating and cooling modes (Digital Pixel Cell Viability and Microscopy Solutions, Science Park Square, Brighton, UK) to maintain a constant temperature.

We used the BioFlux 200 system (Fluxion Biosciences) for all microfluidic experiments, with the exception of the microfluidic experiments shown in Fig. 3 (see below). When using the BioFlux 200 system, we followed closely our previously published experimental protocol (23). Briefly, exponential phase cells were introduced in dual-inlet glass bottom microfluidic channels (24-well plate, product-number 910-0050) and allowed to attach to the surface in the absence of flow for 10-15 min. After flushing out the planktonic and weakly adhered cells from the test section, we kept the flow rate constant at 4 µl h^−1^, (or 12 µl h^−1^ in the high flow speed experiments, Fig. 2*C* and *D*), and imaged cells using brightfield microscopy at a rate of 1 frame min^−1^. Stable antibiotic gradients were generated in the test section of the microfluidic channels via molecular diffusion by injecting growth medium through both inlets and the test antibiotic through just one of them (Fig. 1).

To estimate the distribution of antibiotic in the dual-inlet BioFlux microfluidic experiments (Fig. 1*A*), we flowed fluorescein through one inlet and buffer through the other. This fluorescent dye was imaged using the aforementioned Zeiss microscope with scanning laser confocal attachment (Zeiss LSM 700), a Zeiss EC Plan Neofluar 10X objective, and a Zeiss LSM T-PMT unit to simultaneously visualize the channel geometry. We recorded z-stacks of images at multiple, overlapping positions along the length of the device and the maximum fluorescent intensity in *z* was used to obtain the relative fluorescein concentration at each *x, y* position (Fig. 1*A*). To simulate the distribution of antibiotics in our BioFlux device (Fig. 1*A* and 2*C*), we developed a mathematical model that incorporates the combined effect of fluid advection and molecular diffusion (see section below for details).

We used epi-fluorescent microscopy to visualise both propidium iodide and the mNeonGreen fluorescent protein of our pyocin reporter strain (Fig. 4*B-F*). Propidium iodide (30 µM) was added to the growth medium that was injected through both inlets of our devices. Epi-fluorescent images were recorded at a frame rate of 0.1 frame min^−1^ with exposure times of 90 and 200 nm for the propidium iodide and mNeonGreen pyocin reporter, respectively.

In the experiments where we analysed the long-term viability of cells that undergo chemotaxis towards antibiotics (Fig. 3), we used fluid-walled microfluidics (36). Unlike the BioFlux assays, where cells are fully surrounded by PDMS and glass walls, these open microfluidic devices allowed us to reconfigure channels mid-experiment to isolate and extract cells as they undergo chemotaxis. These experimental methods are explained in detail in a companion publication (37). Briefly, fluid-walled channels were fabricated using Dulbecco’s Modified Eagle Medium (DMEM) + 10% Fetal Bovine Serum (FBS) on untreated, 50 mm glass-bottomed culture dishes (MatTek) using a custom-designed ‘printer’ (iotaSciences Ltd, Oxford, UK). Interfacial forces pin the infused media to the glass surface, which is then overlaid with the fluorocarbon FC40 (iotaSciences Ltd), an immiscible and bio-compatible liquid that prevents evaporation of the fluid within the channels. TB growth medium was then infused through the device to replace the DMEM + FBS used for fabrication (36).

Antibiotic gradients were generated using ‘m-shaped’ devices where the flow from two inlet arms merges together in a central channel (37). Using the printer, a 1:1 co-culture of YFP-labelled WT and unlabelled *ΔpilG* cells was infused into the device at 0.4 µl min^−1^ and the cells were then left to attach to the bottom glass surface of the channel for 10 min in the absence of flow. The device was then transferred to the microscope and growth media was infused into the two inlets using 25G needles at a rate of 0.1 µl min^−1^ per inlet. The needles where connected to 500 µl glass syringes (Hamilton) that were mounted on separate syringe pumps (PhD Ultra, Harvard Apparatus) using 1 m of tubing (PTFE, Adhesive Dispensing Ltd). Ciprofloxacin (100X MIC) was added to one of the syringes to generate a steady gradient in the channel (37). The growth media in both syringes was supplemented with 10% spent medium in order to elicit the chemotactic response more rapidly (see Fig. 2*B*). To ensure the flow was steady over the course of the experiment, fluid was withdrawn at the same rate it was injected (0.2 µl min^−1^) from a circular sink at the end of the test channel using a third syringe (Plastipak, 10 mL, Becton Dickinson) mounted on a third syringe pump (PhD Ultra, Harvard Apparatus). Cells were imaged using brightfield microscopy immediately downstream of the junction between the two inlet arms at a rate of 1 frame min^−1^. Epifluorescence imaging was used to compare the distribution of unlabelled *ΔpilG* and YFP-labelled WT cells at the beginning and end of the experiment.

After a sufficient number of cells had migrated towards ciprofloxacin (≈24 h), we re-configured the central channel of the device using a PTFE (polytetrafluoroethylene) stylus (38) mounted to the microscope condenser to isolate different populations of migrating cells within a separate fluid chamber (Fig. 3*A*-*D*; see (37) for details). Immediately prior to this re-configuration, we reduced the concentration of ciprofloxacin flowing through one of the inlet arms from 100X MIC to 3X MIC using a fourth syringe (Plastipak, 1 mL, Becton Dickinson) mounted on a separate syringe pump (PhD Ultra, Harvard Apparatus). During this process, there was a ≈20 s period during which flow passed through the 100X MIC and the 3X MIC syringes simultaneously and this shifted the antibiotic gradient towards the antibiotic-free side of the channel. In this period, any cells that had only recently initiated chemotaxis would therefore experience a very brief increase in background antibiotic concentration.

We also expect that the channel reconfiguration process might generate lateral flows that could potentially transport a small number of antibiotic naïve-cells into the isolated fluid chamber. To ensure that none of these antibiotic naïve-cells were able to grow, cells within the fluid chambers were exposed to 3X MIC ciprofloxacin for 2 h. In separate experiments that exposed antibiotic-naive cells to 3X MIC ciprofloxacin in liquid culture, fewer than 1 in 10^5^ cells survived after 2 h. Even in the experiment where the channel was reconfigured at Position A (the position for which the largest number of cells were isolated, Fig. 3*A*), we estimated that there were only approximately 1000 cells in the fluid chamber. Therefore, even if all 1000 cells in the chamber were antibiotic-naive, we would expect that 99% of the time all cells would be killed by the subsequent treatment. This, along with our finding that only cells isolated from a single experiment were found to be viable, suggests that contamination of the isolated fluid chambers with antibiotic naïve-cells was not an issue.

After 2 h, the contents of the isolated fluid-chambers were extracted and replaced five times with antibiotic-free growth medium (TB) using the printer, to ensure minimal ciprofloxacin remained within the chambers. During each of the five extractions, the contents were spread onto separate antibiotic-free LB agar (1.5%) plates and monitored for colony growth (Fig. 3*F*). We also imaged the entire bottom surface of the isolated chamber in brightfield every 30 min to monitor the viability of the surface-attached cells that remained within the chambers (Fig. 3*G*). To confirm that all of the antibiotic had been removed from the chambers, ≈25 exponential-phase, antibiotic-naive cells were added at the end of each experiment, which were observed subsequently to undergo rapid cell division, covering the entire bottom surface of the chamber within approximately 24 h (Fig. 3*G*).

### Study of secretions and other factors produced by cells

To test the effect of diffusible cell products on twitching chemotaxis we followed our previously published protocol (13). Briefly, exponential phase cells were inoculated in 6-well plates in 4 ml of tryptone broth (TB) at a starting optical density (at 600 nm) of 0.25. Cultures in these 6-well plates were then allowed to grow at 22°C for 20 h under static conditions in order to obtain a dense culture in which cell-produced factors are present at relatively high concentrations (13). Cultures were centrifuged at 3000 g for 10 min and then filter-sterilised using 0.22 μm filters (Millipore) to obtain cell-free supernatant, which was stored at 4°C. Fresh cell-free supernatant was prepared on the day of each of the microfluidic experiments.

To obtain cell-free supernatant from cells growing under flow conditions within our microfluidic channels (Fig. S6), we used the BioFlux 200 system and followed the same protocol outlined above, except that instead of using dual-inlet microfluidic channels, we used single-inlet channels (48-well plate, product-number 910-0004) where 24 experiments can be run in parallel on the same well plate. By comparison, only 8 experiments can be run in parallel with dual-inlet devices. Flowing tryptone broth through 24 channels simultaneously allowed us to collect a relatively large (≈4 ml) volume of spent growth medium from the outlet wells at the end of the 16 h long experiment. As described above for cultures growing under static conditions, this spent media was then centrifuged at 3000 g for 10 min and filter-sterilised using 0.22 μm filters (Millipore) to obtain cell-free supernatant.

### Image analysis

All images were processed using the open source software Fiji (50) and its associated plugins, and all tracking data was exported from Fiji to Matlab (Mathworks) for subsequent analysis as described previously (23).

We quantified chemotaxis by calculating the movement bias (*β*), which is defined as the number of cells moving up the gradient divided by the number of cells moving down the gradient. Because twitching motility is inherently jerky due to the stochastic dynamics of individual pili (51), we used the direction of a cell’s net displacement over the entire length of its trajectory in our calculation of *β* (rather than the direction of its instantaneous displacement) and considered only trajectories that were at least 1 h in duration. In addition, we excluded non-motile cells, cell clusters and cells that do not exhibit appreciable movement from their starting position from these analyses. To do this, we calculated a cell’s net to gross displacement ratio (NGDR) – the straight-line distance between a trajectory’s start and end positions divided by the total distance travelled by a cell in that time – and excluded any cells with an NGDR of less than 0.15 (23). However, for the experiments that used 100% cell-free supernatant collected from cells growing under flow conditions (Fig. S6), cells reached high densities relatively quickly and so trajectories were likely to be shorter and more frequently broken by neighbouring cells. To account for these differences, we included all trajectories that were over 15 min in duration and we excluded cells that had an NGDR of less than 0.2. Finally, in all of our analyses, we restricted our measurement of movement bias to cells in the central region of the device, where the chemical gradients were largest (*G* = 1/*C*_MAX_ ∂*C/*∂*y* > 0.001, Fig. 2*C*). While our measurements of the movement bias considered only the motile cells in the central region of our microfluidic device, we note that each of our analyses is typically based on thousands of individual cell trajectories.

To quantify how the number of cells attached to the surface changes over time in our isolated fluid chambers (Fig. 3*G*), we used a timeseries of brightfield images collected every 30 min. These images were processed first using the “Normalise Local Contrast” and then the “Subtract Background” plugins in Fiji to remove variations in the intensity of the background. Images were then thresholded to obtain a binary image of the cells. The fraction of the surface covered by cells was then measured by calculating the number of pixels corresponding to cells divided by the total number of pixels.

Brightfield and fluorescent images were processed in a similar manner in order to compare the numbers of YFP-labelled WT and unlabelled *ΔpilG* cells remaining in isolated chambers following channel reconfiguration (Fig. 3*E*). The total number of cells in a given region was calculated by using our previously outlined cell tracking pipeline to count the number of cells present in our brightfield images (23). This number was then compared to the number of cells in our fluorescent images to calculate the relative proportion of WT and *ΔpilG* cells, pooling data across four different bio-replicates.

A similar image processing pipeline was used to quantify how pyocin expression varies across the width of the microfluidic device (Fig. 4*D*-*F*). We again detected the pixels corresponding to cells by thresholding brightfield images and then we thresholded the epifluorescence images that quantified pyocin expression. Using these two binary images, we calculated the proportion of pixels within cells that also corresponded to appreciable pyocin expression. This quantity estimates the relative proportion of cells that are expressing pyocin and was binned in rows of 10 pixels along *y* to calculate how pyocin expression varies across the width of the device. To reduce the effect of solitary cells, we smoothed this proportion with a moving average filter.

We used previously described methods (23) to both automatically detect when twitching cells reverse direction and quantify how the reversal rate differs for cells traveling up the gradient versus down the gradient (Fig. S4). Reversals are relatively rare events (cells reverse direction approximately once every several hours) so a relatively large number of cells must be tracked to obtain reliable measurements of the reversal rate. Cells were imaged at a frame rate of 1 min for ≈10 h and at each time point, a cell can either carry on moving in a relatively straight line or reverse direction. Given our large sample size and the low probability that a cell reverses direction at any given time point, we assumed that reversals can be modelled using a Poisson distribution, which we use to calculate the confidence intervals of our experimentally measured reversal rates.

The rose plots shown in Fig. 1 show histograms of the overall trajectory movement direction, measured from a trajectory’s origin to its terminus (calculated using the ‘ksdensity’ function in Matlab, MathWorks), for all trajectories longer than 10 min.

### Modelling the distribution of antibiotics and the distribution of a hypothetical antibiotic-induced cell product within the microfluidic device

We used a mathematical model of diffusion to simulate the distribution of antibiotic within our devices (or analogously the distribution of cell products originating upstream of the test section in the absence of an antibiotic). We use this to illustrate how this distribution would differ from that of hypothetical cell products generated in response to antibiotic within the test section of the device (antibiotic-induced cell products).

The distribution of antibiotics in our device is expected to closely follow the distribution of a gradient of fluorescein, which has a similar diffusion coefficient and is readily imaged using confocal fluorescence microscopy (Fig. 1*A*, (23)). A previous study found that this fluorescein gradient can be accurately modelled by solving the time dependent diffusion equation in one spatial dimension (23), which has an analytical solution (52). Briefly, this model assumes the flow through the test section of the device moves at a constant speed, so one can estimate how long the two streams of fluid from each inlet have been in contact with one another at each *x* position along the device, allowing the time dependent solution to be transformed into a spatial map of the steady-state concentration field at each [*x*, *y*] position in the device (23). Here we assumed no-flux boundaries at *y* = −175 and 175 µm and an “initial condition” such that the normalised antibiotic concentration is unity from *y* = −175 µm to 0 and zero from *y* = 0 to 175 µm at *x* = 0. This model allowed us to predict the location of the 1X MIC isocontour under both high and low flow conditions, which are in good agreement with the distribution of biofilm experimentally observed in our microfluidic device (Fig. 2*C*). This model was also used to group cell trajectories into different bins according to the strength of the antibiotic gradient they have been exposed to (Fig. 2*D*). Here, the flow rate is 1.1 × 10^6^ µm^3^ s^−1^ (“low flow speed”) or 3.3 × 10^6^ µm^3^ s^−1^ (“high flow speed”) and the diffusion coefficient is assumed to be 200 µm^2^ s^−1^, which is a typical value for low molecular weight compounds dissolved in water.

We used a similar modelling framework to simulate the distribution of a hypothetical cell product produced only in the presence of the antibiotic. In the test section of our dual-inlet microfluidic experiments, cells exposed to sub-MIC concentrations of antibiotic formed biofilms (Figs. 1*A*, 2*C*), where they could hypothetically generate compounds in response to the antibiotic. Intuitively, one would expect that the concentration of such a compound would increase as fluid moves in the downstream direction, as it passes over the top of more and more antibiotic-exposed cells. To model this scenario, we again used a time dependent, one dimensional model of diffusion, assuming that the concentration of any antibiotic-induced cell product is zero at *x* = 0 (the “initial condition”). This assumption is supported by our observation that cells upstream of the test section are rapidly cleared from the arm of the device containing antibiotics, suggesting that the amount of antibiotic-induced cell products originating upstream of the test section was negligible.

Whilst it is difficult to speculate how cells exposed to different concentrations of antibiotic might differ in the rate at which they produce an antibiotic induced product, for simplicity we assumed that cells on the top half of the device generate this compound at a constant rate per unit area. More specifically, we included a source term in our model of diffusion that is equal to unity from *y* = 0 to 175 µm and zero for *y* = −175 to 0 µm. We numerically integrated the governing diffusion equation using the “pdepe” function in Matlab (Mathworks). For consistency, we assumed that the diffusion coefficient of the hypothetical antibiotic-induced cell product is the same as that of the antibiotic (200 µm^2^ s^−1^), but the general trends illustrated in Fig. S8 do not depend on the exact value of *D*.

The distribution of antibiotic and the distribution of the hypothetical antibiotic-induced cell product show starkly different patterns. The strength of the antibiotic gradient is maximum when the fluid from the two inlets first meet one another at *x* = 0 and rapidly decreases further downstream. In contrast, the strength of the hypothetical antibiotic-induced cell product is zero at *x* = 0 and increases steadily as one moves downstream (Fig. S8). Our massively parallel cell tracking analyses show that the movement bias is predicted by the former distribution (Fig. 2*D*), suggesting the directed movement of cells in our experiments is not driven by a cell product induced by a sub-MIC concentration of antibiotics.

## Acknowledgments

We thank Joanne Engel and Yuki Inclan for the Δ*pilG* mutant, plasmid and complemented mutant strains, along with Michael Hopkins and Diego Gonzalez for the Δ*pilG* and respective wild-type chromosomally tagged with constitutively-expressed yellow fluorescent proteins. We thank Leslie Vanderpant (Digital Pixel Cell Viability and Microscopy Solutions, Brighton, UK) for the custom microscope incubation chambers used in this study. We also thank Alvaro San Millán and Craig MacLean for advice. This work was funded by Fundação para a Ciência e Tecnologia (SFRH-BD-73470-2010), BBSRC (BBSRC Discovery Fellowship, BB/T009098/1) and Wellcome Trust (Junior Interdisciplinary Fellowship) to NMO; the Human Frontier Science Program (LT001181/2011L and RGY0080/2021), EPSRC Pump Priming Award (EP/M027430/1) and BBSRC New Investigator Grant (BB/R018383/1) to WMD; and by European Research Council Grant 787932 and Wellcome Trust Investigator award 209397/Z/17/Z to KRF.

## Supplemental Information

**Figure S1.**
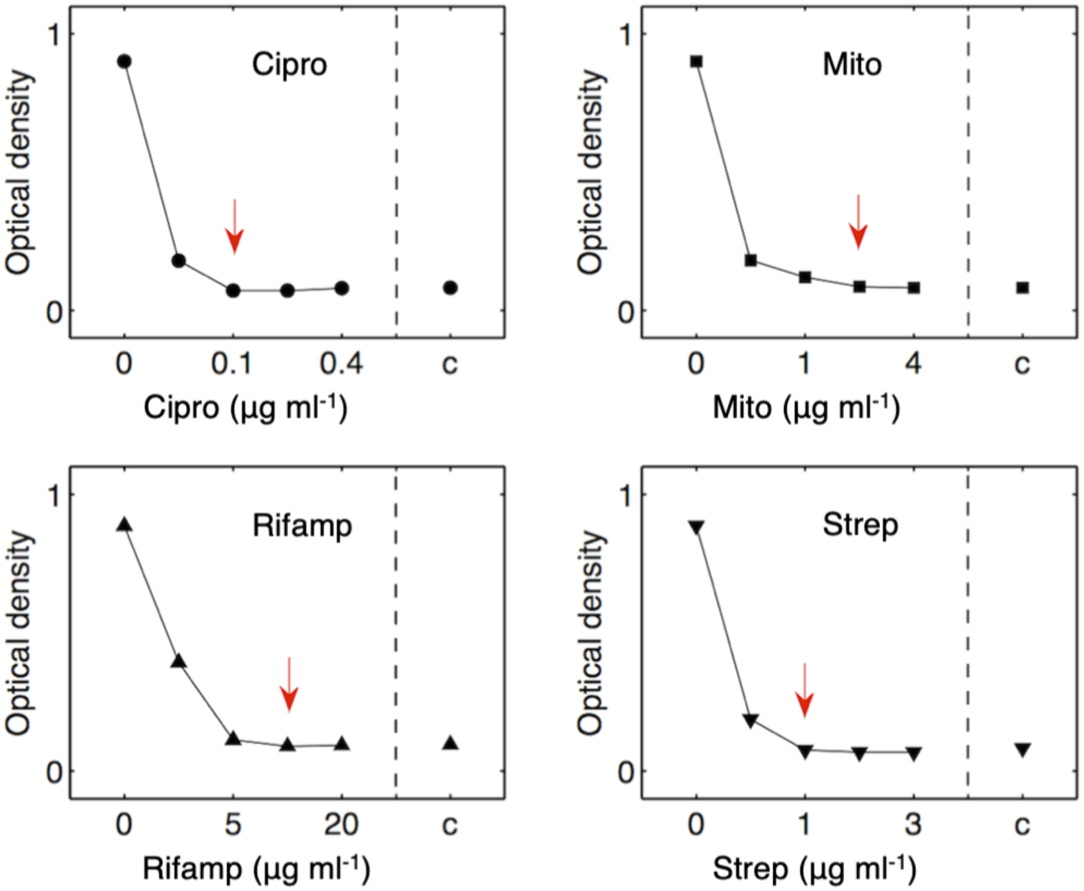
Quantifying the minimum inhibitory concentration (MIC) of different antibiotics in shaken culture. Here we show the optical density after 20 h in different concentrations of ‘Cipro’ = ciprofloxacin, ‘Mito’ = Mitomycin C, ‘Rifamp’ = Rifampicin, and ‘Strep’ = Streptomycin. ‘c’ denotes controls without cells conducted in pure tryptone broth. Red arrows denote the minimum inhibitory concentration (MIC) for each antibiotic, which is defined here as the minimum concentration of antibiotic that prevents detectable growth after 20 h. Error bars representing SE of eight independent bio-replicates are too small to see and fall within each marker shown.

**Figure S2.**
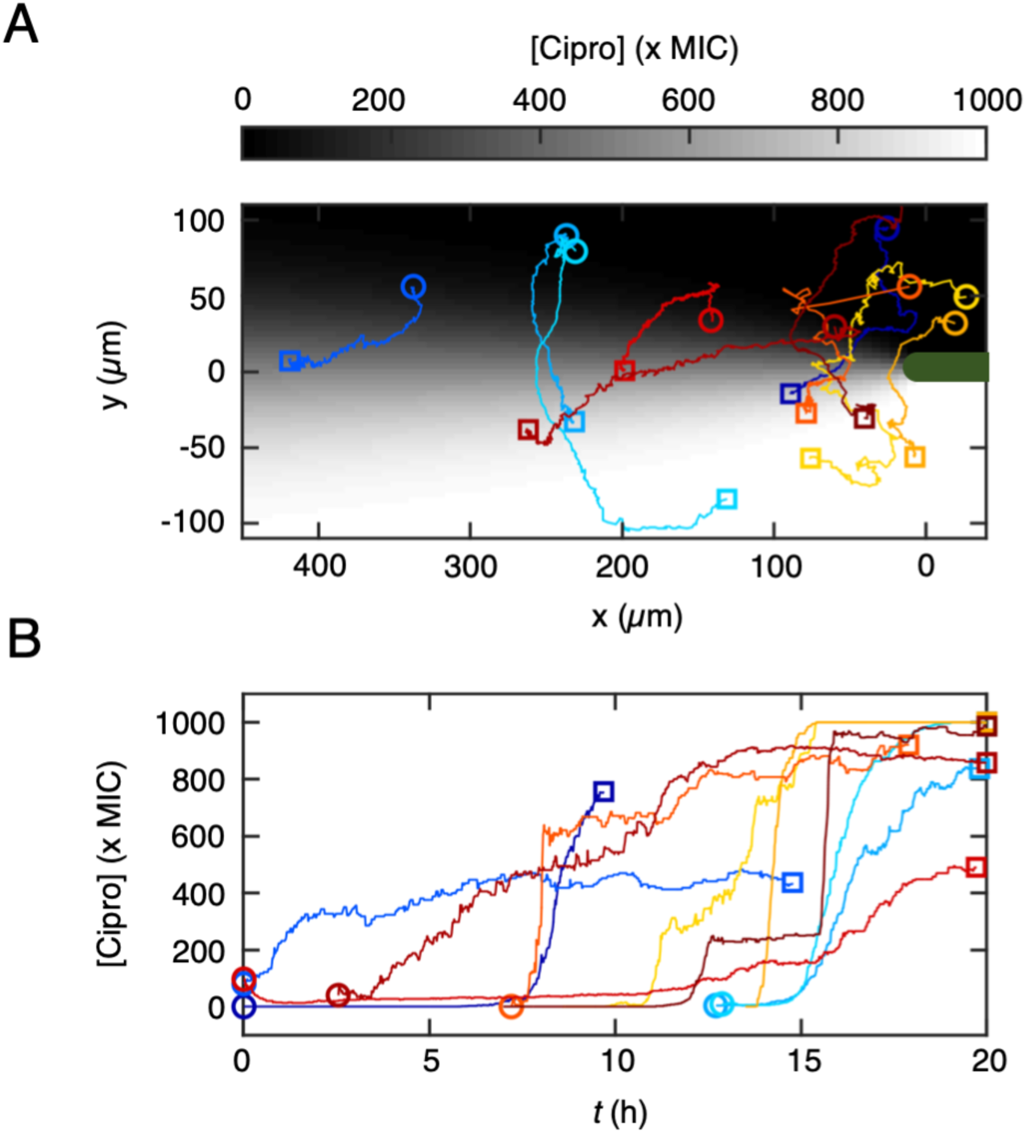
Cells that chemotax towards ciprofloxacin remain motile for hours after reaching antibiotic concentrations corresponding to hundreds of times that of their MIC. **(A)** While automated tracking can simultaneously follow the motility of hundreds of cells, cell trajectories periodically break when two cells come into close proximity and touch one another. To quantify cell movement over long periods of time in the densely crowded conditions found in our experiments, we therefore used manual cell tracking to follow the movement of 10 representative cells moving in a ciprofloxacin gradient that varies from a concentration of zero to 1000 times the MIC. **(B)** We used the mathematical model of ciprofloxacin distribution within our device (Materials and Methods) to resolve the concentration that these cells experienced over time. This shows that cells are capable of moving into concentrations hundreds of times that of the MIC and remain motile for hours, even though they are ultimately non-viable (Fig. 3).

**Figure S3.**
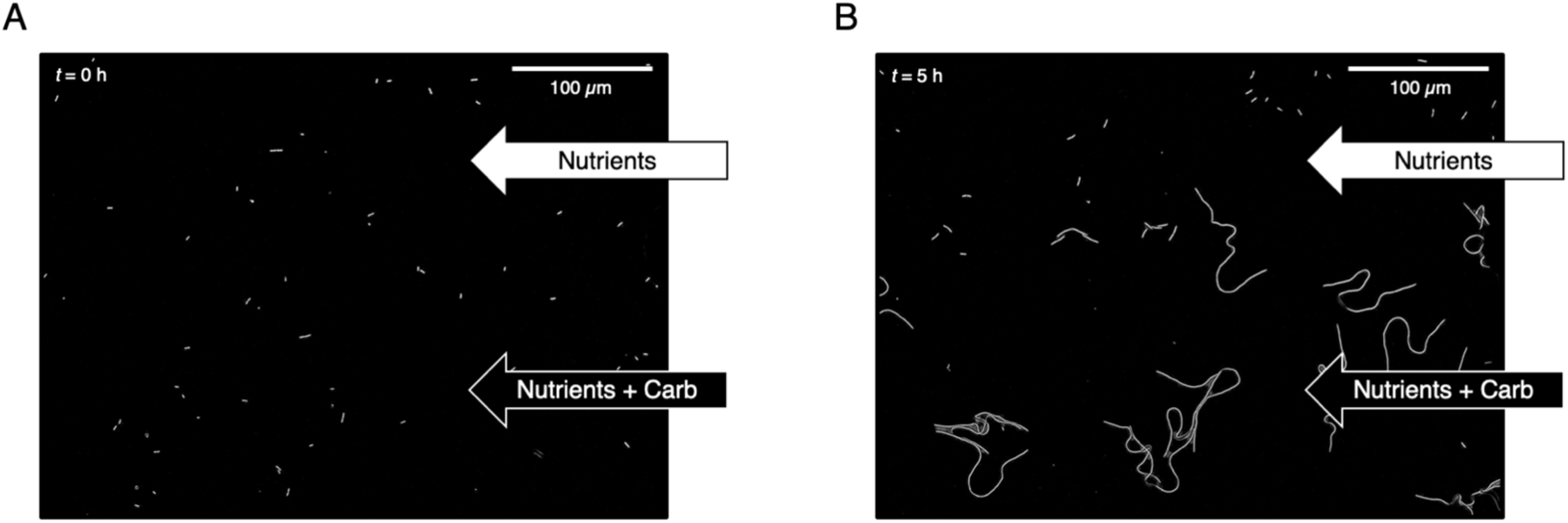
Carbenicillin causes extreme cell elongation, preventing cell motility. **(A, B)** We exposed cells to a gradient of carbenicillin *(C*_MAX_ = 10X MIC) to test whether *P. aeruginosa* also undergoes chemotaxis towards *β*-lactam antibiotics. After five hours of exposure to carbenicillin, cells become highly elongated and lack the ability to move.

**Figure S4.**
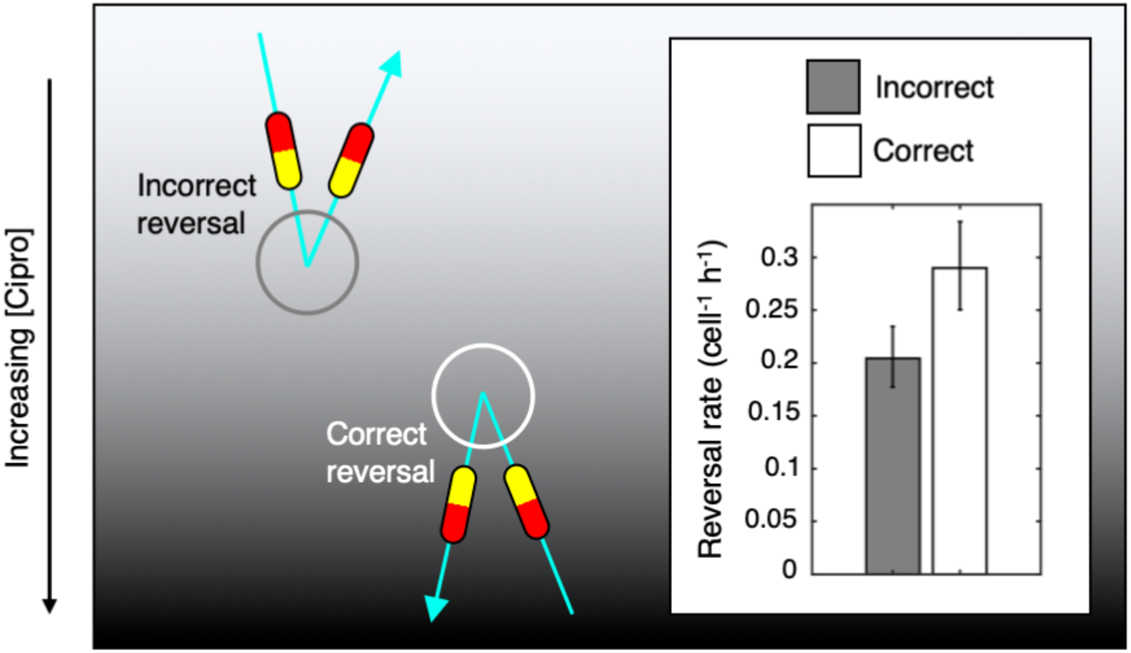
*P. aeruginosa* biases its motility towards ciprofloxacin by actively reversing direction. Twitching *P. aeruginosa* **cells** have the ability to actively regulate the direction of their movement by reversing direction approximately 180° (23, 53). In the context of a chemoattractant gradient, these reversals can either be “correct” (white circle), which reorient a cell previously moving away from the chemoattractant, or “incorrect” (grey circle), which do the opposite. Whilst reversals are relatively rare (i.e. approximately once every five hours), automated cell tracking allows us to follow the movement of thousands of cells and automatically identify when reversals occur (23). In ciprofloxacin gradients, we find that cells are significantly more likely to undergo correct reversals (white bar, inset) than incorrect reversals, which is consistent with previous observations of chemotaxis (23). Error bars show 95% confidence intervals, assuming that reversals follow a Poisson distribution (Materials and Methods).

**Figure S5.**
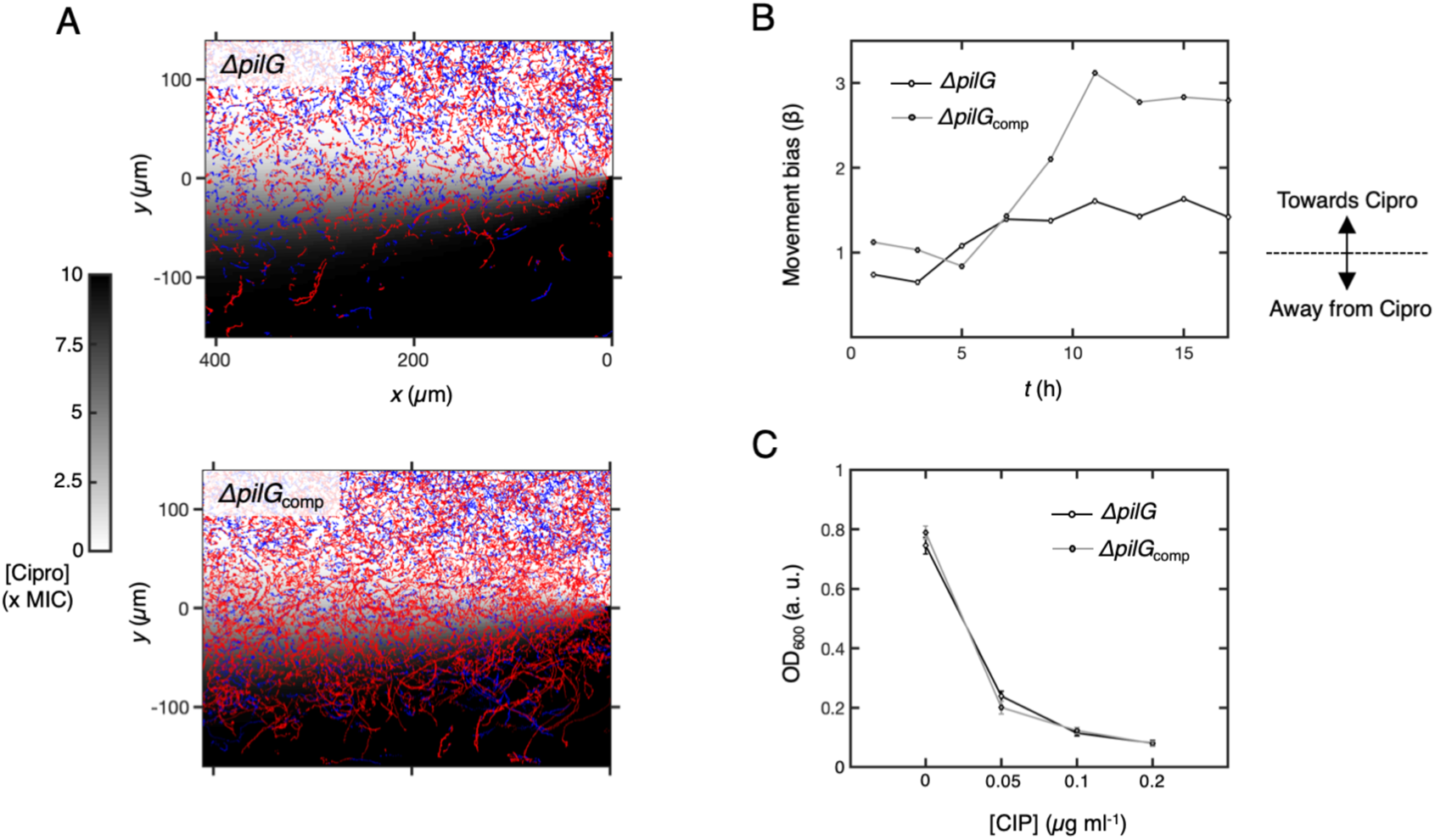
A *ΔpilG* strain that cannot reverse direction does not bias its movement towards ciprofloxacin. **(A)** To test the role of reversals in driving chemotaxis towards antibiotics, we studied a *ΔpilG* mutant that has previously been shown to have a reduced reversal rate and is unable to perform chemotaxis (23). We exposed both the mutant strain (*ΔpilG*) and its complement (*ΔpilG*_comp_) to a gradient of ciprofloxacin (*C*_MAX_ = 10X MIC ciprofloxacin). Cell trajectories are colour coded by whether they move towards increasing (red) or decreasing (blue) ciprofloxacin concentrations. **(B)** Whilst the movement of *ΔpilG* cells remains unbiased throughout the experiment (*β* ≈ 1), the *ΔpilG*_comp_ strain begins to move towards increasing ciprofloxacin after ≈7.5 h (*β* > 1). **(C)** The optical density of *ΔpilG* and *ΔpilG*_comp_ cells in different ciprofloxacin concentrations are nearly the same after an incubation of 20 h in shaken culture, and are similar to previous estimates of PAO1 ((27), Fig. S1), suggesting that PilG does not appreciably affect ciprofloxacin resistance. As noted in the main text, *ΔpilG* cells exhibit reduced motility compared to WT cells in these assays (23) and so our analyses only include cells of each genotype that make appreciable movement from their initial position (Materials and Methods).

**Figure S6.**
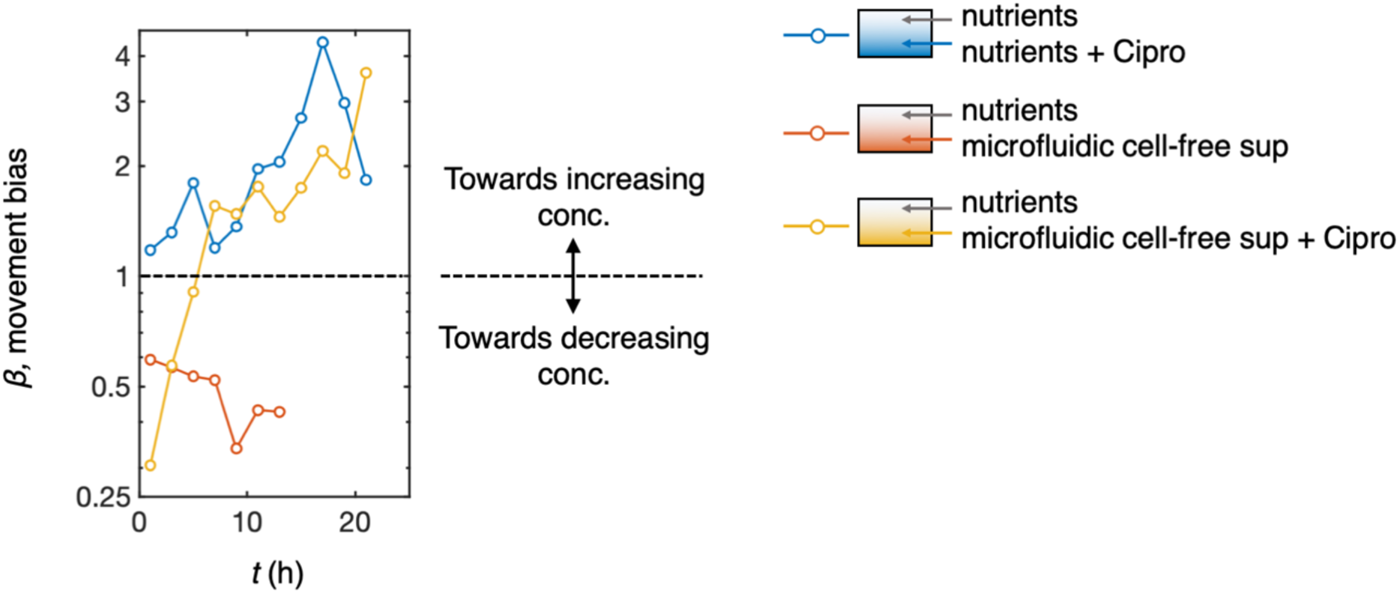
Cells move towards antibiotics despite repulsion from cell-free supernatant collected from microfluidic devices. Cells bias their movement away from 100% cell-free supernatant obtained from microfluidic devices (red line, *β* < 1), as they do for cell-free supernatant collected from static-well plates (Fig. 2B). Note that after ≈13 h, the cell density in this gradient, (in which there are no antibiotics present), becomes too high to track cells reliably. If ciprofloxacin (*C*_MAX_ = 10X MIC) is added to the supernatant (yellow line), cells initially move away from the antibiotic (*β* < 1), but after ≈7 h cells begin moving towards the added ciprofloxacin (*β* > 1). A control experiment with a ciprofloxacin gradient only (*C*_MAX_ = 10X MIC) is shown for comparison (blue line). See also Movie 3.

**Figure S7.**
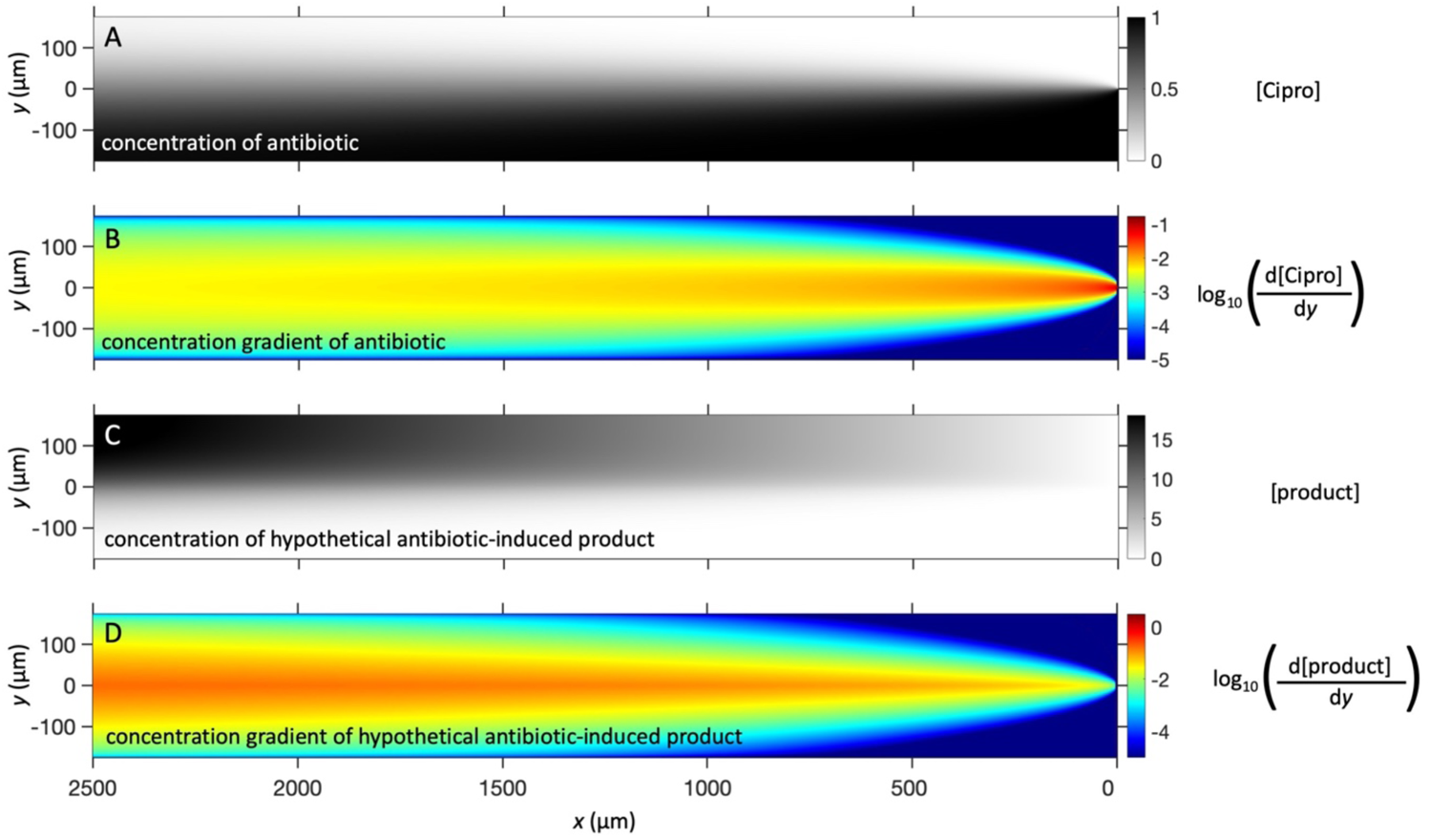
A hypothetical cell product released from cells in response to antibiotics would increase in the downstream direction of our device, producing a starkly different distribution compared to that of the antibiotic itself. Here we used a mathematical model of diffusion to simulate both the distribution of antibiotic within the microfluidic device (A and B), and the distribution of a hypothetical product that is released by cells in response to the antibiotic, which here is assumed to be produced at a constant rate in the region *x* = 0 to 175 µm (C and D). The concentration and concentration gradients have been normalised and are presented here in arbitrary units. See Materials and Methods for further details.

**Figure S8.**
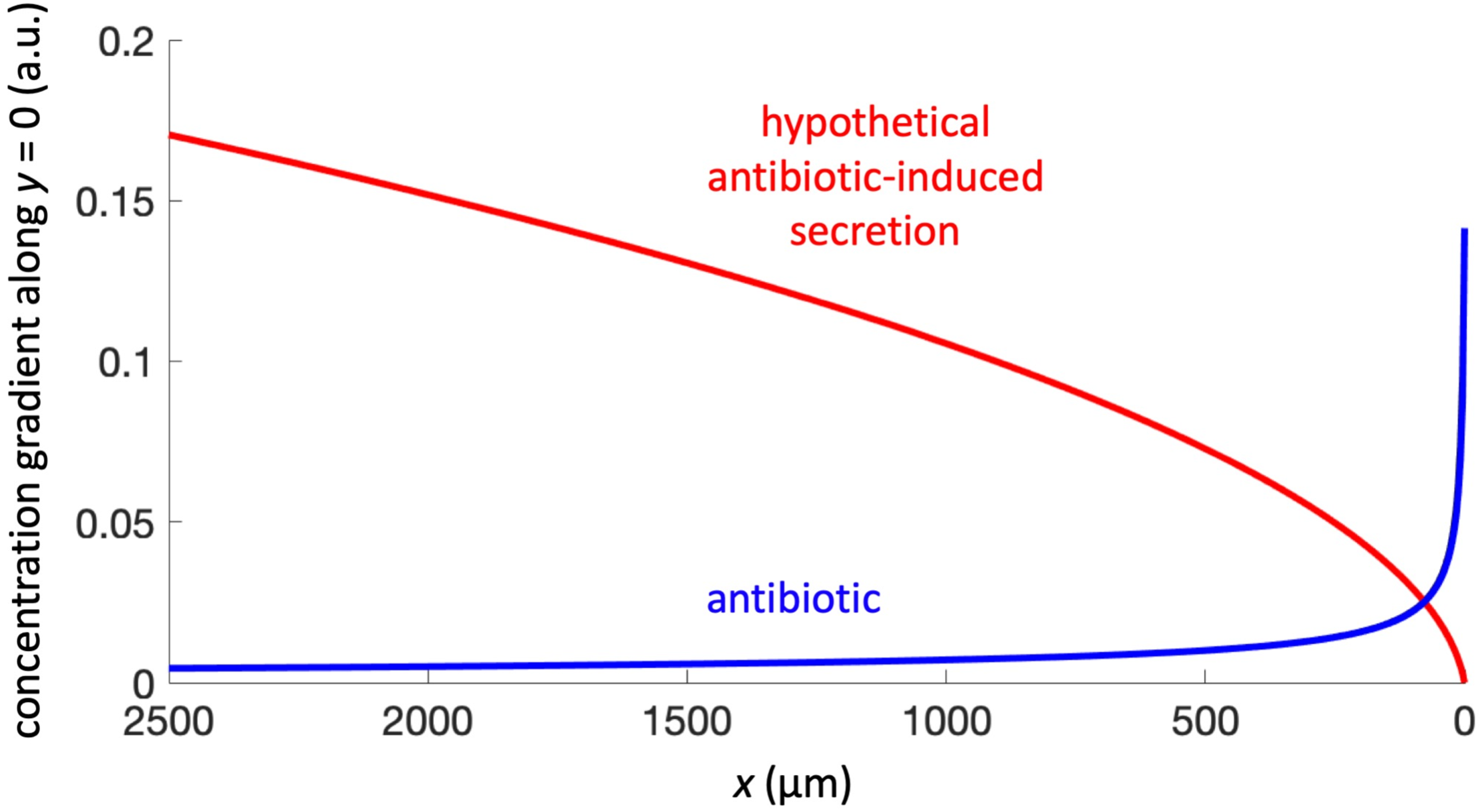
Predictions of chemical gradients in the microfluidic device. The concentration gradient of the antibiotic is predicted to decrease in the downstream direction, whilst the concentration gradient of a hypothetical antibiotic-induced cell product would increase in the downstream direction. The blue and red lines show model predictions of d[antibiotic]/d*y* and d[product]/d*y*, respectively along the centreline of the channel (*y =* 0, see Fig. S7).

