## Supplementary material for "Suicidal chemotaxis in bacteria": Movie Legends

**Movie 1.** Suicidal chemotaxis towards antibiotics. Surface-attached *P. aeruginosa* cells move via twitching motility towards the antibiotic ciprofloxacin ($C_{MAX}$ = 10X MIC), after an initial period of random motility. A static gradient of ciprofloxacin is generated using a dual-inlet microfluidic device with nutrients (tryptone broth) flowing through one inlet and nutrients supplemented with ciprofloxacin through the other. The panel on the right-hand side shows a magnified view of the region marked with the dashed white box in the first frame.

**Movie 2.** A reversal-null *ΔpilG* mutant does not chemotax towards antibiotics. Cells were exposed to a static gradient of ciprofloxacin ($C_{MAX}$ = 10X MIC), with nutrients (tryptone broth) flowing through the top-half of the channel and nutrients supplemented with ciprofloxacin flowing through the bottom-half of the channel. The right-hand panel shows *ΔpilG* mutant cells, which are unable to reverse direction and bias their motility towards ciprofloxacin (Fig. S5). In contrast, the left-hand panel shows complemented mutant cells ($\Delta pilG_{comp}$), whose ability to undergo chemotaxis towards ciprofloxacin is restored. Note that *ΔpilG* cells exhibit reduced motility compared to WT cells in these assays (23) and so we only analyse cells that show appreciable movement from their initial position (Materials and Methods).

**Movie 3.** The effects of cell-free supernatant on chemotaxis with and without antibiotics. Left-hand side shows chemotaxis towards antibiotics, where surface-attached *P. aeruginosa* cells move via twitching motility towards ciprofloxacin ($C_{MAX}$ = 10X MIC; see also Movie 1). Middle panel shows cells in a gradient of cell-free supernatant ($C_{MAX}$ = 100% cell free supernatant) extracted from cells grown in tryptone broth within microfluidic devices (‘microfluidic cell-free sup’). Cells are repelled by the cell-free supernatant. Right-hand side shows cells in a combined gradient of cell-free supernatant *and* ciprofloxacin. Cells move towards the ciprofloxacin despite the repulsion of cell-free supernatant (shown in the middle panel).

**Movie 4.** Cells release bacteriocins (pyocins) during chemotaxis towards antibiotics (ciprofloxacin, $C_{MAX}$ = 10X MIC). Cells turn green when they make bacteriocins as they have a mNeonGreen fluorescent transcriptional reporter for Pyocin R2. Pyocins are released via programmed cell lysis, which can be seen in the movie where it is followed by a transient burst of red colour. This colour is caused by the presence of the dye propidium iodide which binds to the DNA released when cells lyse.
